# SOD1 activity thresholds and TOR signalling modulate VAP(P58S) aggregation via ROS-induced proteasomal degradation in a *Drosophila* model of Amyotrophic Lateral Sclerosis

**DOI:** 10.1101/368100

**Authors:** Kriti Chaplot, Lokesh Pimpale, Balaji Ramalingam, Senthilkumar Deivasigamani, Siddhesh S. Kamat, Girish S. Ratnaparkhi

## Abstract

Familial Amyotrophic Lateral Sclerosis (F-ALS) is an incurable, late onset motor neuron disease, linked strongly to various causative genetic loci. *ALS8* codes for a missense mutation, P56S, in VAMP-associated Protein B (VAPB) that causes the protein to misfold and form cellular aggregates. Uncovering genes and mechanisms that affect aggregation dynamics would greatly help increase our understanding of the disease and lead to potential therapeutics.

Here, we develop a quantitative high-throughput, *Drosophila* S2R+ cell-based kinetic assay coupled with fluorescent microscopy to score for genes involved in the modulation of aggregates of fly ortholog, VAP(P58S), tagged with GFP. As proof of principle, we conducted a targeted RNAi screen against 900 genes, consisting of VAP genetic interactors, other ALS loci, as also genes involved in proteostasis. The screen identified 150 hits that modify aggregation, including the ALS loci *SOD1, TDP43* and also genes belonging to the TOR pathway.

To validate these modifiers, we developed a system to measure the extent of VAP(P58S) aggregation in the *Drosophila* third instar larval brain using the *UAS-GAL4* system, followed by quantitative imaging of cellular inclusions. We find that reduction of SOD1 activity or decreased TOR signalling reduces aggregation. Interestingly, we find that increase in cellular reactive oxygen species (ROS) levels, assessed by measuring oxidation of cellular lipids and proteins, in response to SOD1 knockdown or by inhibition of TOR signalling appears to be the trigger for clearing of aggregates. The mechanism of aggregate clearance is, primarily, the proteasomal machinery, and not autophagy. Increase in VAP, but not VAP(P58S) levels, appears to elevate ROS, which may in turn regulate *VAP* transcription in a feedback loop.

We have thus uncovered an interesting interplay between SOD1, ROS and TOR signalling that regulates the dynamics of VAP aggregation. Mechanistic processes underlying such cellular regulatory networks will lead us to a better understanding of initiation and progression of ALS.

## Introduction

Amyotrophic lateral sclerosis (ALS) is a progressive, fatal neurodegenerative disease characterized by loss of motor neurons resulting in muscular atrophy, gradual paralysis and ultimately death of the patient within 2-5 years post diagnosis (Cleveland and Rothstein 2001; Tarasiuk *et al*. 2012). Most often, the disease occurs sporadically (S-ALS). However, in ∼10% of the cases, the disease occurs due to inheritance of altered gene(s) (F-ALS). *ALS1/SOD1* coding for superoxide dismutase 1, was the first causative locus to be discovered (Deng *et al*. 1993; Rosen *et al*. 1993), with more than 170 SOD1 mutations attributed to the diseased state. Since then, about 50 potential genetic loci (Taylor *et al*. 2016) have been identified in ALS through Genome-wide association (GWAS), linkage and sequencing studies. Recent studies have emphasized on the oligogenic basis for ALS (Van Blitterswijk *et al*. 2012; Deivasigamani *et al*. 2014), suggesting that ALS loci may be a part of a gene regulatory network (GRN) that breaks down late in the life of a diseased individual. At the cellular level, several hallmarks of ALS include breakdown of cellular homeostasis (Cluskey and Ramsden 2001), endoplasmic reticulum (ER) stress, unfolded protein response, aggregation, oxidative stress, mitochondrial dysfunction and autophagy. While several studies have demonstrated the wide-range of consequences of the genetic alterations on cellular function, no clear unifying mechanism has emerged that might explain the pathogenesis of the disease (Andersen and Al-Chalabi 2011; Walker and Atkin 2011; Mulligan and Chakrabartty 2013; Turner *et al*. 2013; Taylor *et al*. 2016).

In 2004, Mayana Zatz’s group (Nishimura *et al*. 2004) discovered a novel causative genetic locus, VAMP-associated protein B (VAPB), termed as ALS8, in a large Brazilian family whose members succumbed to ALS and/or Spinal muscular atrophy (SMA). The point mutation of P56S was identified in the N-terminal, Major Sperm Domain (MSP) of VAPB (Nishimura *et al*. 2004). VAPB is an integral membrane protein present in the ER membrane, ER-Golgi intermediate compartment, mitochondrial-associated membrane and the plasma membrane, implicated in important functions in the cell such as vesicular trafficking, ER structure maintenance, lipid biosynthesis, microtubule organization, mitochondrial mobility and calcium homeostasis (Lev *et al*. 2008; Murphy and Levine 2016). Recent studies have highlighted its critical role in membrane contact sites (Alpy *et al*. 2013; Gomez-Suaga *et al*. 2017b; Metz *et al*. 2017; Yadav *et al*. 2018; Zhao *et al*. 2018). The *Drosophila* ortholog of VAPB is VAP33A/CG5014 (Called VAP hereafter) and has been used to develop models for ALS (Chai *et al*. 2008; Ratnaparkhi *et al*. 2008; Deivasigamani *et al*. 2014; Moustaqim-Barrette *et al*. 2014; Sanhueza *et al*. 2015). We have previously identified a *Drosophila* VAP GRN comprising of 406 genes, including a novel interaction with the mTOR pathway (Deivasigamani *et al*. 2014). The ALS8 mutation can also alter VAP’s physical interaction with other proteins, including FFAT motif containing proteins (Loewen *et al*. 2003; Murphy and Levine 2016), impairing cellular functions (De Vos *et al*. 2012; Moustaqim-Barrette *et al*. 2014; Huttlin *et al*. 2015). Ubiquitinated cellular aggregates (Ratnaparkhi *et al*. 2008; Papiani *et al*. 2012) are seen on VAP mutant expression, and are capable of sequestering the wildtype VAP protein in a dominant negative manner (Teuling *et al*. 2007; Ratnaparkhi *et al*. 2008). In *Drosophila*, neuronal overexpression of VAP(P58S), and subsequent formation of aggregates, in the background of endogenous VAP appear to lead to only mild neurodegenerative phenotypes, such as flight defects, as compared to the more severe phenotypes associated with wild type VAP neuronal overexpression (Ratnaparkhi *et al*. 2008; Tsuda *et al*. 2008). Previously, we have used the UAS-GAL4 system to study the interaction between VAP and mTOR signalling using the NMJ phenotype associated with neuronally overexpressed VAP(P58S)(Deivasigamani *et al*. 2014). The functional consequence of neuronal VAP(P58S) aggregation in this system is not fully understood and its contribution to disease remains elusive.

In this study, we identify 150 genetic modifiers of VAP(P58S) *aggregation* by conducting a directed S2R+ cell based RNAi screen, targeting 900 unique genes belonging to different categories that are associated either with ALS or VAP function or proteostasis. We used the previously described (C155-Gal4;UAS-VAP(P58S)) system (Ratnaparkhi *et al*. 2008; Deivasigamani *et al*. 2014) to validate one such modifier, SOD1, *in vivo*, in the third instar larval brain of *Drosophila* by measuring changes in aggregation of VAP(P58S) in response to modulation of *SOD1* levels. Our data indicates that clearance of VAP(P58S) aggregates via the proteasomal machinery is enhanced by inducing reactive oxygen species (ROS) due to loss of SOD1 function. We also find a similar clearance of aggregation, attributed to proteasomal degradation, with mTOR downregulation accompanied by elevated ROS. We find that wild type VAP, but not mutant VAP, elevates ROS. Accumulated ROS results in inhibition of endogenous *VAP* transcription, a phenomenon that may directly affect both familial as well as sporadic ALS pathogenesis.

## Results

### A Drosophila S2R+ cell culture model to study VAP(P58S) aggregation

C-terminal and N-terminal fusions of VAP and VAP(P58S) with GFP were used to transfect cells and generate stable S2R+ lines, as described in Materials & Methods (Fig. 1A, Suppl. Fig. 1A). VAP:GFP showed a non-nuclear, reticular localization in the cell with <10% of the transfected (GFP-positive) cells showing high intensity puncta (Fig. 1B, Suppl. Fig. 1A). In contrast, >80% of the GFP-positive VAP(P58S):GFP, cells showed distinct high intensity puncta with little or no background staining within the cell (Fig. 1C, Suppl. Fig. 1A). Super resolution imaging confirmed that VAP appeared to be reticular, while VAP(P58S) was found in inclusion bodies (Fig. 1D). In contrast, GFP, when expressed showed a uniform cytoplasmic signal (Suppl. Fig. 1B). Both N-terminal GFP fusions, GFP:VAP and GFP:VAP(P58S) showed puncta formation at levels comparable to VAP(P58S):GFP, and hence were not used further in the study (Suppl. Fig. 1A). All further experiments (see next section) were carried out with stable lines expressing VAP:GFP or VAP(P58S):GFP, which showed expected/relevant localization and levels of aggregation.

**Figure 1:**
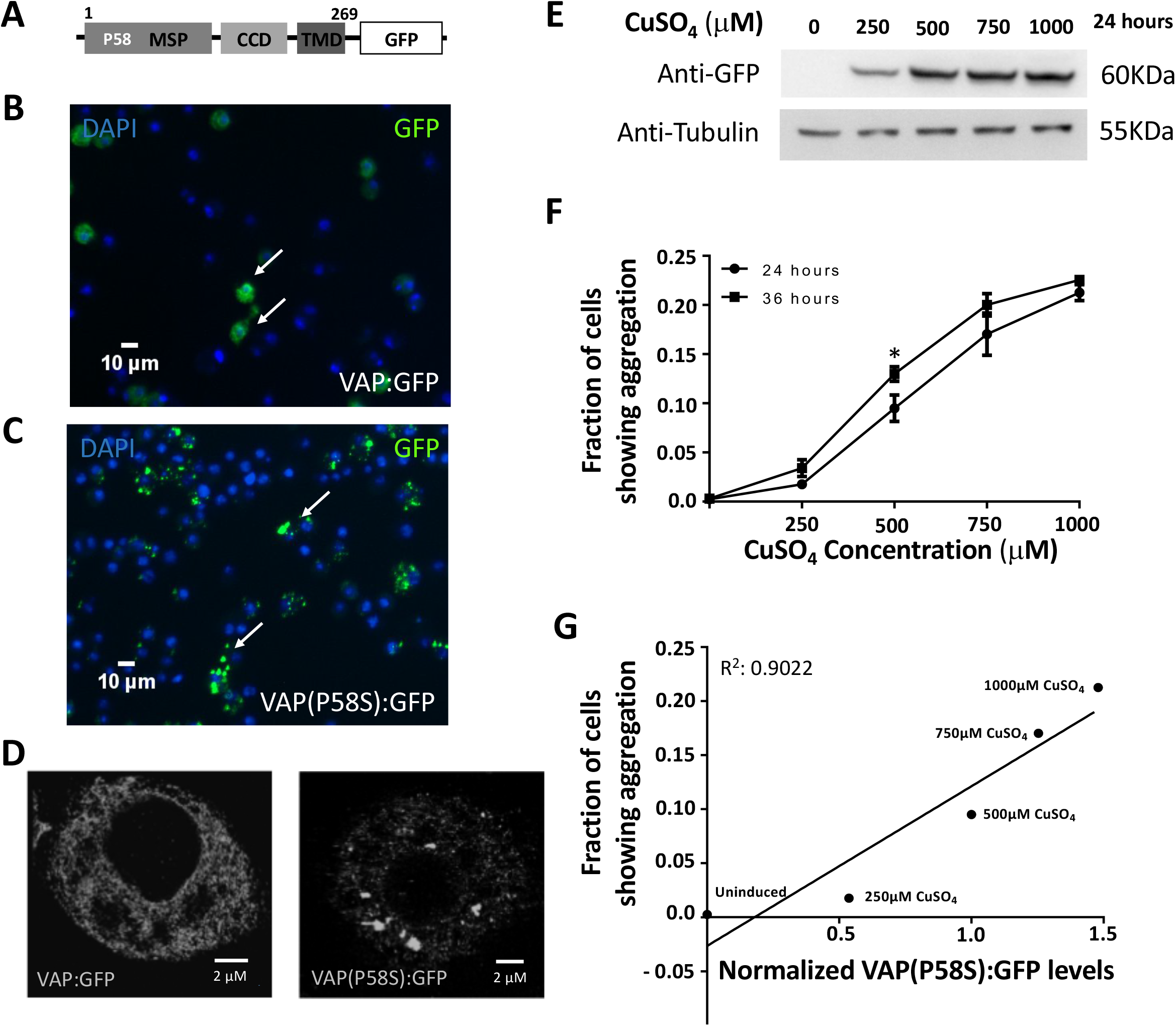
A *Drosophila* cell culture model to study VAP(P58S) aggregation. **A**. VAP:GFP and VAP(P58S):GFP when expressed in S2R+ cells allow efficient visualization of VAP protein in the cell by epifluorescence. **B, C**. Stable cell lines: expressing *VAP(P58S):GFP*, under an inducible metallothionein promoter, results in aggregation (C), unlike VAP:GFP wild-type (B). GFP is visualized by epifluorescence and chromatin by DAPI, postfixation. **D**. A super resolution image, using Ground State Depletion microscopy, showing GFP inclusions forming in cells expressing VAP(P58S):GFP but not in VAP:GFP. **E**. VAP(P58S):GFP protein levels in cells increase with increasing CuSO_4_ concentration at 24 hours post induction. **F**. Increase in fraction of cells showing GFP positive inclusions increases with CuSO_4_ concentration. At 500 µM CuSO_**4**_, inclusions significantly increase between 24 hours and 36 hours. Student’s t-test (P-value: *<0.01) **G**. A linear correlation between fraction of cells showing aggregation, measured using microscopy plotted against relative VAP(P58S):GFP protein levels, as quantified by western blotting, at 24 hours post induction.

### An S2R+ cell based reverse genetics screen is developed to identify modifiers of VAP(P58S) aggregation

In an attempt to identify genetic modifiers of VAP(P58S) aggregation kinetics, we conducted a focused S2R+ cell based RNAi screen, targeting 900 unique genes belonging to nine different categories or families associated with ALS or VAP function. We generated stable S2R+ cell lines expressing VAP(P58S):GFP under a Cu^2+^ induced promoter. The inducible cell culture system allowed us to increase the VAP(P58S):GFP protein levels in the cell with increasing copper sulphate (CuSO_4_) concentrations (250μM, 500μM, 750μM and 1000μM) at 24 hours post induction (Fig. 1E). Using fluorescence microscopy, we found a linear relationship between the copper sulphate (CuSO_4_) concentrations and also the fraction of cells showing VAP(P58S):GFP aggregates that also increased with time (24 and 36 hours) post induction (Fig. 1F). The concentration dependent increase in relative levels of VAP(P58S):GFP correlated with an increase in fraction of cells showing aggregates (Fig. 1G), indicating the propensity of the mutant protein to aggregate. Early time points (12–16 hours) gave very few cells with aggregates; while non-linearity, high confluency, and cell death became a concern at time points beyond 48 hours and concentrations greater than 750 µM. The aggregation kinetics curve was used to define the extent of aggregation in the cell culture system and select optimum parameters to conduct the RNAi screen. Keeping a modest confluency and well-separated cells for ease of imaging, the screen was performed at a fixed concentration of 500 µM CuSO_4_ at 24 and 36 hours post induction.

We chose 900 genes (Suppl. Table 1A), based on their availability in the Open Biosystems Library (See Materials & Methods) to screen for modifiers that could change aggregation levels of VAP(P58S):GFP. A Gene Ontology (GO) chart (Fig. 2A) represents the biological process associated with these 900 genes, as defined by FlyBase. The genes were selected and categorized (Suppl. Table 1B) on the following basis. First, known *Drosophila* Orthologs of ALS loci (20 genes) and ALS related genes (36 genes) as tabulated in the online ALS database (ALSOD) were chosen. The next category included 273 genes from a VAP *Drosophila* GRN comprising of 406 genes (Deivasigamani *et al*. 2014). As *mTOR* was identified as a major interactor of *VAP* in our previous study (Deivasigamani *et al*. 2014), we chose 22 genes of the extended mTOR pathway. To explore the functional aspects of VAP(P58S), we also screened genes involved in lipid biosynthesis (92 genes) and FFAT motif interactors of VAP (34 genes). In order to identify a role of proteostasis in aggregation, we screened genes involved in unfolded protein response (123 genes), ubiquitin–proteasomal pathway (212 genes), and autophagy (88 genes)

**Figure 2:**
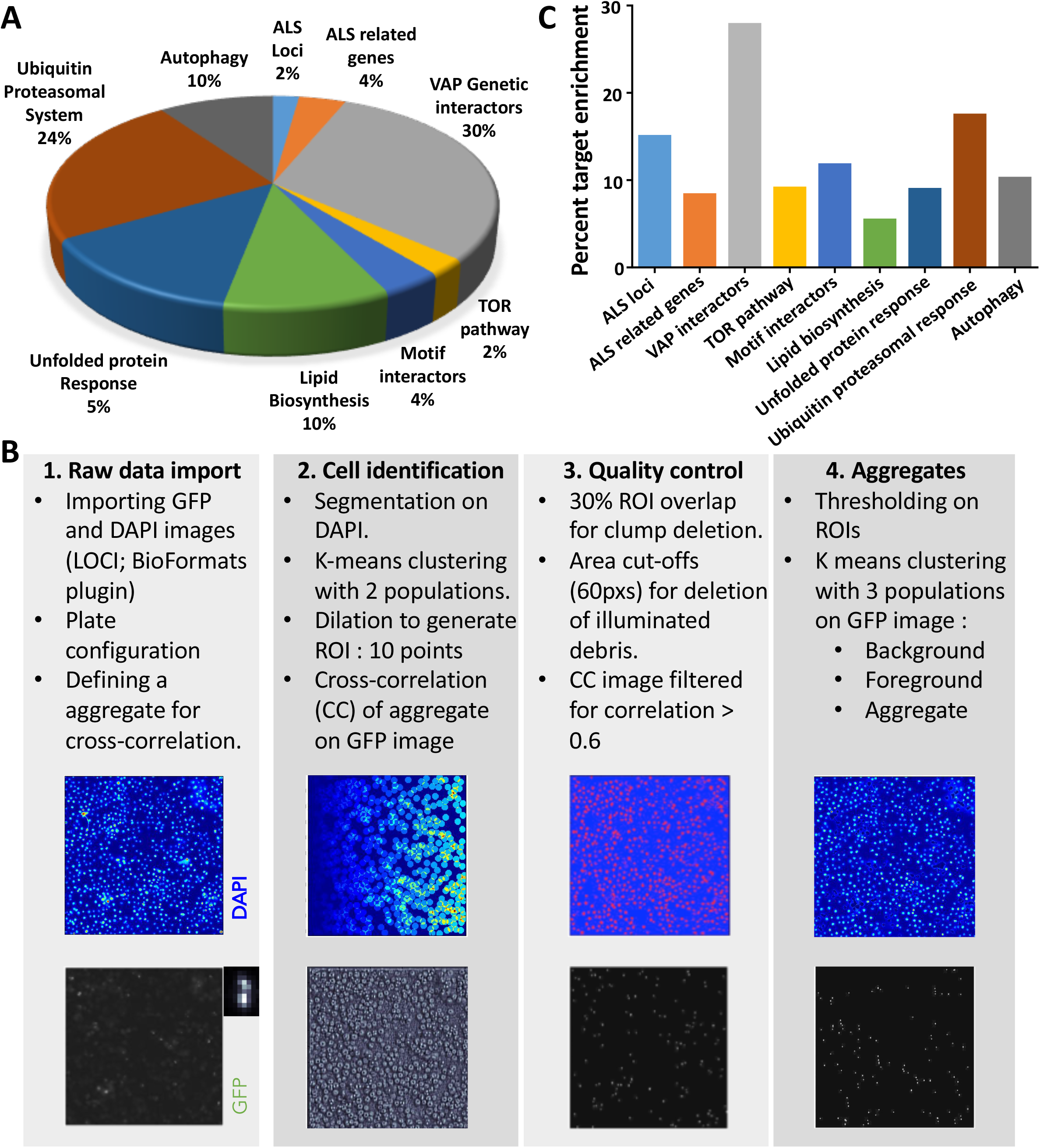
A targeted dsRNA screen in S2R+ cells to discover modifiers of VAP(P58S):GFP aggregation. **A:** dsRNA for 900 genes (Suppl. Table 1A) were chosen for knockdown. GO representation indicates the categories of genes chosen and fraction (%) for each category. Genes were categorized as described in text (Supplementary table 1A, 1B). **B:** Workflow of the steps executed for image analysis using an automated MATLAB script (Dey *et al*, 2014). Steps detailed in Material and Methods. **C:** The end result of the screen is a list of 150 genes identified based on average cell intensity, which have been found to modify aggregation of VAP(P58S):GFP. Graph indicates the percent fold enrichment of targets within each gene category. Genes are listed in Supplementary table 1C.

The images collected at the end of the screen (detailed in Materials and Methods) were analysed by an automated MATLAB analysis (see Materials & Methods; Fig. 2B). Based on average cell intensity, 150 targets (Suppl. Table 1C), and based on total cell intensity, 85 targets (Suppl. Table 1D) that modulated VAP(P58S):GFP aggregation kinetics were identified; 55 genes were found to be targets as per both parameters. Enrichment profile of target genes are plotted in Fig 2C and Suppl. Fig. 1C. ALS loci, notably *SOD1* and *TBPH*, were found as interesting modulators perturbing VAP(P58S):GFP aggregation. Targets belonging to the VAP genetic network, as defined by (Deivasigamani *et al*. 2014), were also enriched. As identified earlier (Deivasigamani *et al*. 2014), components of the mTOR pathway also appeared to be key regulators of VAP(P58S):GFP aggregation. However, less than 10% of genes screened belonging to families associated with lipid biosynthesis and motif interactors, were identified as targets, suggesting lower functional relevance for VAP(P58S):GFP. Interestingly, genes related to ubiquitin proteasomal system such as ubiquitin ligases and proteasome components were enriched, as were the autophagy related genes such as *ATG7* and *ATG3*. From the unfolded protein response category, along with chaperones such as heat shock proteins, we also identified peptidyl prolyl isomerases as targets. Overall, in our primary targeted screen, we found various genetic interactors of wildtype VAP as modulators of VAP(P58S) aggregation as well; importantly, the uncovering of two ALS loci, *SOD1* and *TDP-43*, mTOR pathway genes such as *Rheb* and *S6K*, and genes enriched in ubiquitin proteasomal system as modulators of VAP(P58S) aggregation dynamics, lead us to develop an *in vivo* model to validate these genes and to understand mechanisms underlying these interactions in the animal.

### A model system for measuring VAP(P58S) aggregation in the Drosophila larval brain

In order to validate targets from the screen *in vivo*, we used the *UAS-GAL4* system to specifically overexpress wild-type *VAP* or *VAP(P58S)* in the brain using a pan-neuronal driver, *C155 (elav)*(Ratnaparkhi *et al*. 2008; Deivasigamani *et al*. 2014). Based on anti-VAP immunostaining, unlike wild-type VAP (Suppl. Fig. 2A), mutant VAP(P58S) formed distinct cellular puncta and could be used as a model to study aggregation in the animal (Suppl. Fig. 2B-D). These aggregates have been shown to be ubiquitinated and dominant-negative when expressed in muscle (Ratnaparkhi et al, 2008). To develop a methodology for quantitation of aggregates in the brain (described in Materials & Methods), we used temperature as a means to increase GAL4 activity, which would increase VAP(P58S) dosage and possibly, aggregation. An increase in mean VAP(P58S) aggregation density was observed from 18 ºC to 25 ºC, but not significantly between 25 ºC and 28 ºC (Suppl. Fig. 2H). Neuronal knockdown of VAP, using RNAi, in *CI55-GAL4/+; UAS-VAP(P58S)/+* flies, at each temperature (Suppl. Fig. 2E-G), led to a significant decrease in corresponding aggregation density of the ventral nerve cord (Suppl. Fig. 2H). The above experiments suggest that at 25 ºC, we could quantify changes in VAP(P58S) aggregation density in the brain of the larvae, and here onwards, we use this system to further validate modifiers of aggregation identified from the cell-based screen.

### Drosophila SOD1 is a modifier of VAP(P58S) aggregation

*SOD1*, first known ALS locus (Rosen *et al*. 1993), has been implicated in both sporadic as well as familial cases and was our first choice for validation of the S2R+ based screen, in the animal. We previously identified *SOD1* as a genetic interactor of *VAP* in a fly-based reverse genetics screen (Deivasigamani *et al*. 2014). Here, we individually knocked down *SOD1* using three independent RNAi lines in the *CI55-GAL4/+; UAS-VAP(P58S)/+* background and observed a significant decrease in aggregation density in the ventral nerve cord (Fig. 3A, 3B, Suppl. Fig. 3A, 3C, 3D). This three-fold decrease in VAP aggregates was comparable to the reduction seen with VAP RNAi. Likewise, we overexpressed *SOD1* in the *CI55-GAL4/+; UAS-VAP(P58S)/+* background. Here, however, we did not find a significant change in aggregation density (Fig. 3C, 3D Suppl. Fig. 3B, 3C, 3E). Taken together, these results suggest a need for a threshold level of *SOD1* to maintain VAP(P58S) inclusions.

**Figure 3:**
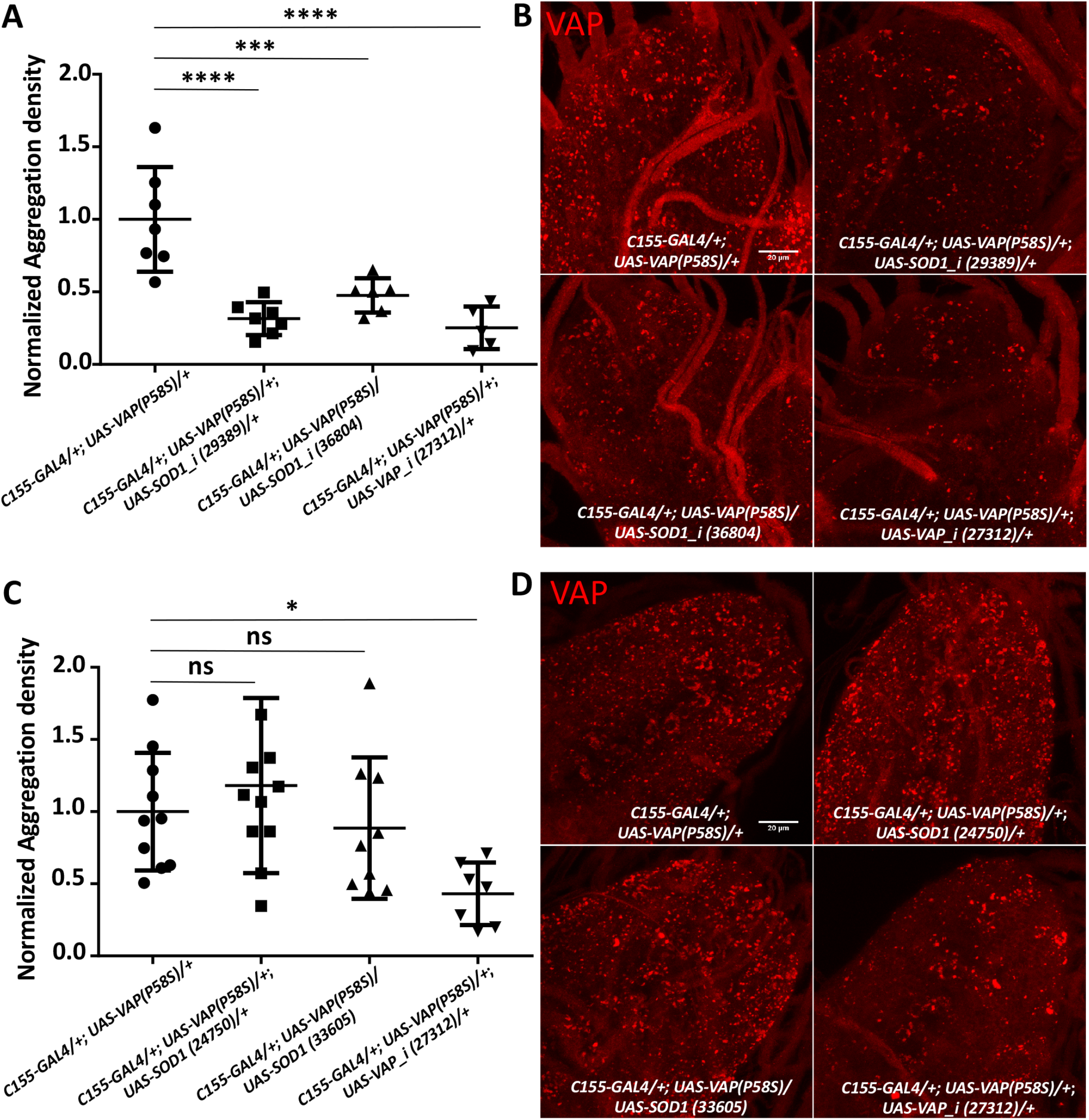
S0D1 loss-of-function reduces VAP(P58S) aggregation in larval brains. **A:** *SOD1* knockdown in the nervous system decreases aggregation density in the ventral nerve cord. VAP knockdown also reduces aggregation due to reduction in VAP protein expression. The ‘_1’ appended to the gene name indicates an RNAi line. ANOVA (P value: ****< 0.0001). Numbers in brackets indicate BDSC stock numbers. **B:** Representative images of the ventral nerve cord showing aggregation of VAP(P58S) with *SOD1* knockdown(29389 and 36804) and with VAP knockdown (27312). **C:** *SOD1* overexpression does not affect aggregation density in the ventral nerve cord. ANOVA (P value: *, 0.0208) **D:** Representative images of the ventral nerve cord showing aggregation of VAP(P58S) with *SOD1* overexpression (24750 and 33605) and with VAP knockdown (27312). All images were taken at the same magnification. Fisher's LSD multiple comparison (P-values, *<0.05, **<0.01, ***<0.001).

### Oxidative stress reduces VAP(P58S) aggregation

Enzymatically, SOD1 metabolizes superoxide species to hydrogen peroxide, thereby preventing oxidative stress. A loss of function of SOD1 would, in principle, increase ROS. We tested whether a chemical mimic, paraquat, which increases cellular ROS (Castello *et al*. 2007; Drechsel and Patel 2008; Cocheme *et al*. 2011), could phenocopy the effect of *SOD1* knockdown. We treated the VAP(P58S):GFP stable line with non-lethal concentrations of 10 mM and 20 mM paraquat for 4 hours prior to CuSO_4_ induction and found that paraquat could significantly reduce the fraction of cells showing GFP positive aggregates (Fig. 4A, Suppl. Fig. 4A) in a dose-dependent manner. Similarly, larvae with the genotype *CI55-GAL4/+; UAS-VAP(P58S)/+* hatched, fed and grown on non-lethal concentration of 5 mM paraquat at 25 ºC, showed a decrease in aggregation density in the third instar larval brain, reminiscent of the *SOD1* knockdown phenotype (Fig. 4B, Suppl. Fig. 4B). We also checked the effect of other ROS scavenging genes such as *SOD2* and *catalase* on VAP(P58S) aggregation. Knockdown of both these genes resulted in a drastic reduction in aggregation density in the ventral nerve cord of *CI55-GAL4/+; UAS-VAP(P58S)/+* larval brains. As seen with SOD1, overexpression of SOD2 did not change aggregation density; however, catalase overexpression resulted in a fractional increase in aggregation density (Suppl. Fig 3F). These results strongly suggest a ROS dependent maintenance and/or stability of VAP(P58S) aggregates.

**Figure 4:**
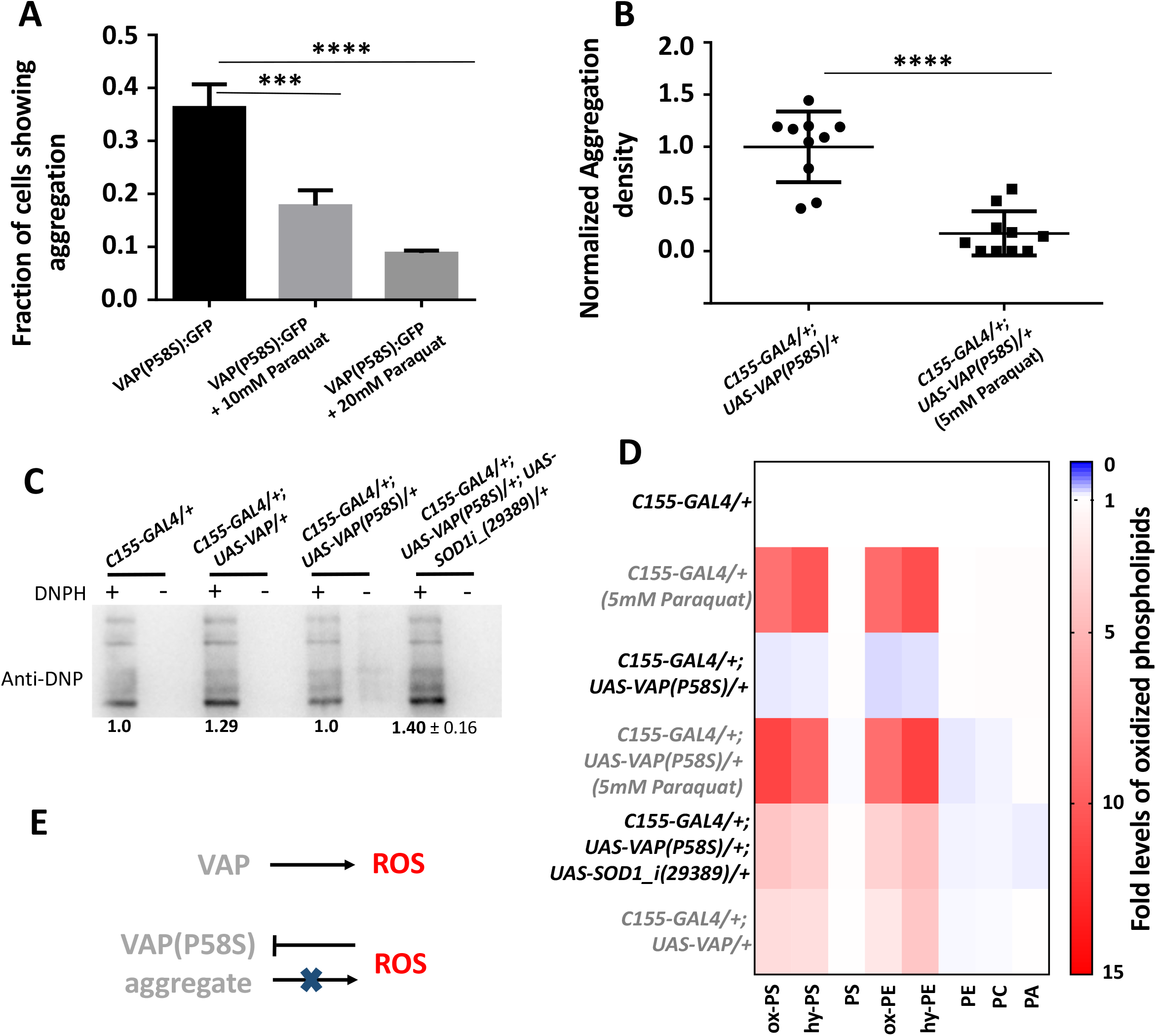
Increase in ROS leads to decrease in VAP(P58S) aggregation levels. **A:** 4 hour Paraquat treatment prior to inducing VAP(P58S):GFP in stable S2R+ cell line, reduces the fraction of cells showing aggregation observed 24 hours post-induction. ANOVA (P-value: ****<0.0001) Fisher’s LSD multiple comparison test (P-values, ***<0.001, ****<0.0001). **B:** Paraquat feeding decreases aggregation density in the ventral nerve cord of third instar larval brains in *C155-GAL4/+; UAS-VAP(P58S)/+* flies. Student’s t-test (P-value: ****<0.0001). **C:** Oxyblot showing increased levels of oxidized proteins in larval brains (N=14) upon SOD knockdown, or VAP overexpression. Values below the gel indicate fold intensity of the strongest band, when compared to control (*C155-GAL4/+)*. *Suppl. Fig. 4C* shows a calibration for the Oxyblot system, values measured after feeding increasing amounts of Paraquat to larvae. **D:** Heat map depicting change in levels of oxidized phospholipids normalized to *C155-GAL4/+*, quantified using MS in response to ROS generated in third instar larval brains (N=4) for the listed genotypes. SOD knockdown as well as VAP overexpression appears to increase cellular ROS levels. Statistical tests are described in *Suppl. table 2*. **E:** Model depicting the effect of overexpression of wildtype and mutant VAP on ROS.

To confirm whether feeding of paraquat and loss of SOD1 function led to an increase in ROS levels in the larval brain, we measured the levels of oxidized proteins and lipids, using the oxyblot kit and quantitative mass spectrometry based lipidomics, respectively. Using the oxyblot assay, we found that feeding *C155-GAL4/+* larvae with increasing concentrations of paraquat (0 mM, 0.05 mM, 0.5 mM, 5 mM) was sufficient to increase ROS in the brain, observed as an increase in intensity of oxidized proteins as compared to unfed larvae (Suppl. Fig. 4C). As expected, neuronal knockdown of *SOD1* in presence of VAP(P58S) aggregates, led to a corresponding increase in intensity of oxidized proteins, demonstrating oxidative stress (Fig. 4C). We found that VAP(P58S) aggregation alone did not significantly change oxidized protein levels as compared to the *C155-GAL4/+* control (Fig. 4C). Unexpectedly, we found that overexpression of *VAP* in neurons led to a distinct increase in oxidation of proteins (Fig. 4C).

To further bolster our findings, we measured levels of oxidized phospholipids in larval brains (Tyurina *et al*. 2000; Kamat *et al*. 2015; Kory *et al*. 2017). On feeding *C155-GAL4/+* larvae with 5 mM paraquat, we enriched and detected 9 oxidized polyunsaturated fatty acids (PUFAs), belonging to phosphatidylserine (PS) and phosphatidylethanolamine (PE) (Fig. 4D, Suppl. Table 2) families of phospholipids, which were significantly elevated in larval brains, compared to the unfed control. PUFA containing oxidatively damaged phospholipids showed a mass addition of +16 (denoted as ox-) or +18 (denoted as hy-) to the parent phospholipid, as a consequence of addition of different ROS. Of note, the parent or precursor phospholipids did not change in concentration, and the concentrations of the oxidized phospholipids were less than 1% of the parent or precursor phospholipids. We found a similar elevation in concentrations of oxidized phospholipids in *C155-GAL4/+; UAS-VAP(P58S)/+; UAS-SOD1_i/+*, but not in *CI55-GAL4/+; UAS-VAP(P58S)/+* which was equivalent to *C155-GAL4/+* control (Fig. 4D, Suppl. Table 2). This elevation in oxidized phospholipids was found to be inversely correlated with corresponding fold change in aggregation density (Suppl. Fig. 4D). Interestingly, we found, as suggested by the oxyblot data, overexpression of *VAP* had a curious effect of increasing oxidation of lipids, indicating that wild type VAP has a cryptic yet important role in regulating ROS levels. Taken together, these results indicate that ROS initiates processes that aid clearance VAP(P58S) aggregates, and is in turn regulated by VAP wildtype levels in the cell.

### ROS activates proteasomal machinery

We further investigated protein degradation mechanisms that may be activated in response to ROS leading to the clearance of VAP(P58S) aggregates. In order to test whether the proteasomal machinery was responsible for reduction in aggregation, we hatched, fed, and grew larvae with proteasomal inhibitor 5µM MG132, and dissected the brains at the wandering third instar stage and analysed the aggregation density. Unfed *C155-GAL4/+; UAS-VAP(P58S)/+; UAS-SOD1_i/+*, as expected, showed reduced aggregation density (Fig. 5C), as compared to unfed control (Fig. 5A, 5E). Upon MG132 feeding, *C155-GAL4/+; UAS-VAP(P58S)/+; UAS-SOD1_i/+*, showed a complete rescue of VAP(P58S) aggregation (Fig. 5D, 5E). Fed *C155-GAL4/+; UAS-VAP(P58S)/+; UAS-SOD1_i/+* also showed an enhanced aggregation density as compared to fed *CI55-GAL4/+; UAS-VAP(P58S)/+*(Fig. 5B, 5E). Aggregates in presence of ROS (with SOD1 knockdown) and proteasomal inhibition (with MG132) appeared to be predominantly smaller, scattered and mislocalized around the nuclear membrane/ER as compared to the respective controls (Fig. 5D’). The localization of the aggregates suggest that may be residing in the Juxta Nuclear Quality Control compartment (JUNQ)-like compartment (Ogrodnik *et al*. 2014). These results indicate that the proteasomal machinery is facilitated in presence of ROS for active degradation of VAP(P58S) aggregates (Fig 5F). However, fed *CI55-GAL4/+; UAS-VAP(P58S)/+* larvae (Fig. 5A) did not show accumulation of aggregation as compared to unfed control (Fig. 5B, 5E), indicating other mechanisms may be at play to maintain the aggregation density.

**Figure 5:**
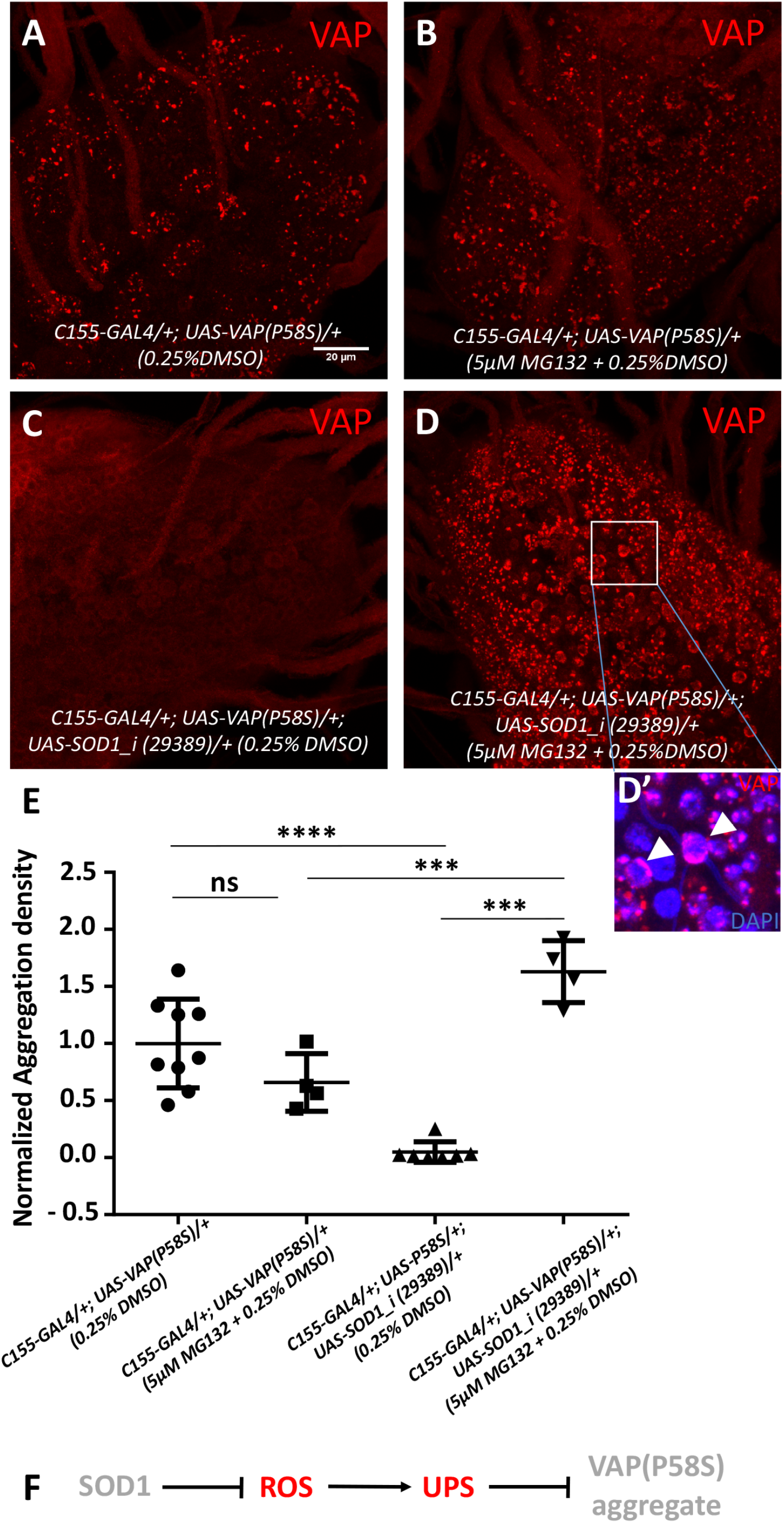
ROS activates proteasomal machinery: **A,B:** MG132 feeding of *C155-GAL4/+; UAS-VAP(P58S)/+*, to inhibit proteasomal machinery, does not accumulate aggregation. **C,D,D’:** MG132 feeding of *C155-GAL4/+; UAS-VAP(P58S)/+; UAS-SOD1_i (29389)/+*, accumulate aggregation. The aggregates, in presence of ROS and MG132, seem to be smaller, scattered and localized around the nuclear membrane (arrowheads) as depicted in inset (**D**’). **E:** Plot showing significant decrease in aggregation density in the ventral nerve cord in *C155-GAL4/+; UAS-VAP(P58S); UAS-SOD1_i (29389)/+ as compared to C155-GAL4/+; UAS-VAP(P58S)/+* control. This decrease is rescued by feeding 5μM MG132 and is significantly higher than the *C155-GAL4/+; UAS-VAP(P58S)/+* control, both unfed and fed with MG132. All images were taken at the same magnification. ANOVA (P-value: ****<0.0001) Fisher's LSD multiple comparison test (P-values, ***<0.001, ****<0.0001) **F:** Model depicting the role of SODl-regulated ROS in activating proteasomal degradation of VAP(P58S) protein/aggregates.

### mTOR inhibition lowers VAP(P58S) aggregation but not via autophagy

We examined whether aggregates could be cleared via autophagy in the third instar larval brain. We inhibited the mTOR pathway by feeding *CI55-GAL4/+; UAS-VAP(P58S)/+* larvae with 200nM rapamycin (Heitman *et al*. 1991) as described (Deivasigamani *et al*. 2014), thereby activating autophagy (Noda and Ohsumi 1998), and observed a drastic clearance of aggregation in the ventral nerve cords as compared to unfed controls (Fig. 6A, 6B, 6C). When *Tor* transcripts were reduced using RNAi in *CI55-GAL4/+; UAS-VAP(P58S)/+*, a similar decrease in aggregation density was found (Fig. 6D, 6E, 6F). However, when autophagy was induced directly via overexpression of Atg1 in *CI55-GAL4/+; UAS-VAP(P58S)/+*, we did not observe clearance of aggregation (Fig. 6G, 6H, 6I). This suggests that mTOR signalling may perturb downstream effectors other than Atg1 which may affect VAP(P58S) aggregation dynamics (Fig 6J).

**Figure 6:**
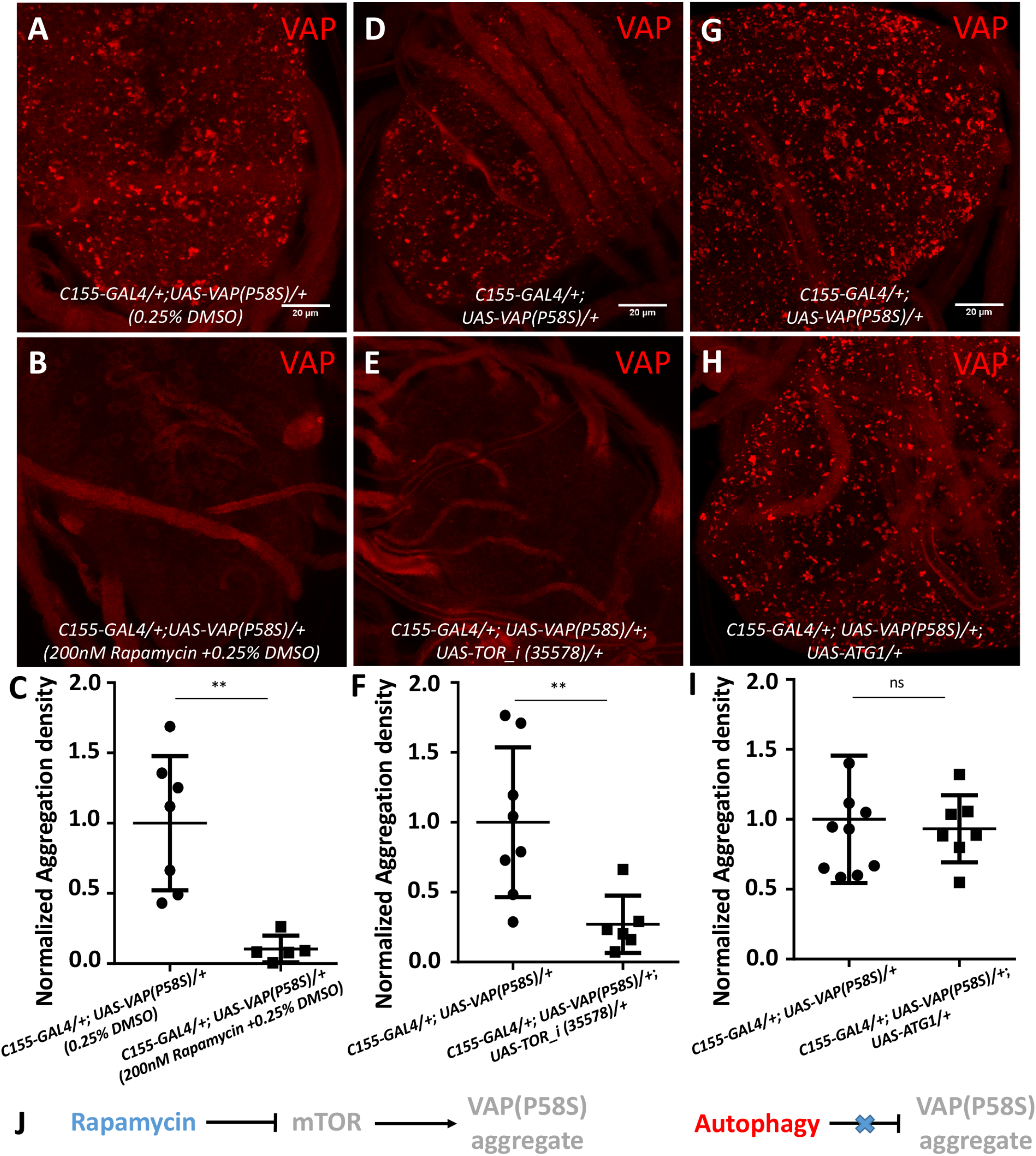
mTOR inhibition reduces VAP(P58S) aggregation independent of autophagy. **A-C:** Rapamycin feeding decreases aggregation density in the ventral nerve cord of third instar larval brains in *C155-GAL4/+; UAS-VAP(P58S)/+* flies. **D-F:** Neuronal *TOR* knockdown decreases aggregation density in the ventral nerve cord. The *‘_i’* appended to the gene name indicates an RNAi line. **G-l:** Neuronal overexpression of Atgl did not affect the aggregation density in the ventral nerve cord. All images were taken at the same magnification. Students’s t-test (P-value, **<0.01) **J:** Model depicting mTOR-regulated clearance of aggregation, independent of autophagy.

### mTOR inhibition promotes proteasomal clearance of VAP(P58S) aggregation via ROS

We first decided to check whether clearance of aggregates with mTOR inhibition correlated with increase in ROS, as in the case of *SOD1* knockdown. We found that levels of several species of oxidized phospholipids were indeed higher with *Tor* knockdown with or without neuronal overexpression of VAP(P58S) in third instar larval brains to levels similar to *SOD1* knockdown (Fig. 7A). mTOR pathway downregulation has recently been shown to activate not only autophagy but also ubiquitin proteasomal machinery (Zhao *et al*. 2015) via Mpk1/ERK5 pathway in yeast and humans (Rousseau and Bertolotti 2016). We tested whether ROS upregulation with *Tor* knockdown could be inducing proteasomal clearance of VAP(P58S) aggregation by feeding *CI55-GAL4/+; UAS-VAP(P58S)/+; UAS-TOR_i/+* with 5µM MG132 (Fig 7B, 7C-E). Although there was a significant decrease in aggregation density with *Tor* knockdown (Fig. 7D), we found only a slight recovery of aggregation in MG132-fed animals (Fig. 7E) as compared to unfed *CI55-GAL4/+; UAS-VAP(P58S)/+* control flies (Fig. 7C). This recovery appeared to be far less dramatic than that seen in the case of *SOD1* knockdown. Taken together, these results indicate that in context of ROS, proteasomal degradation could be the major pathway responsible for clearance of VAP(P58S) aggregation (Fig. 7F), although other downstream effectors of mTOR signalling, including autophagy, cannot be conclusively ruled out as additional mechanisms.

**Figure 7:**
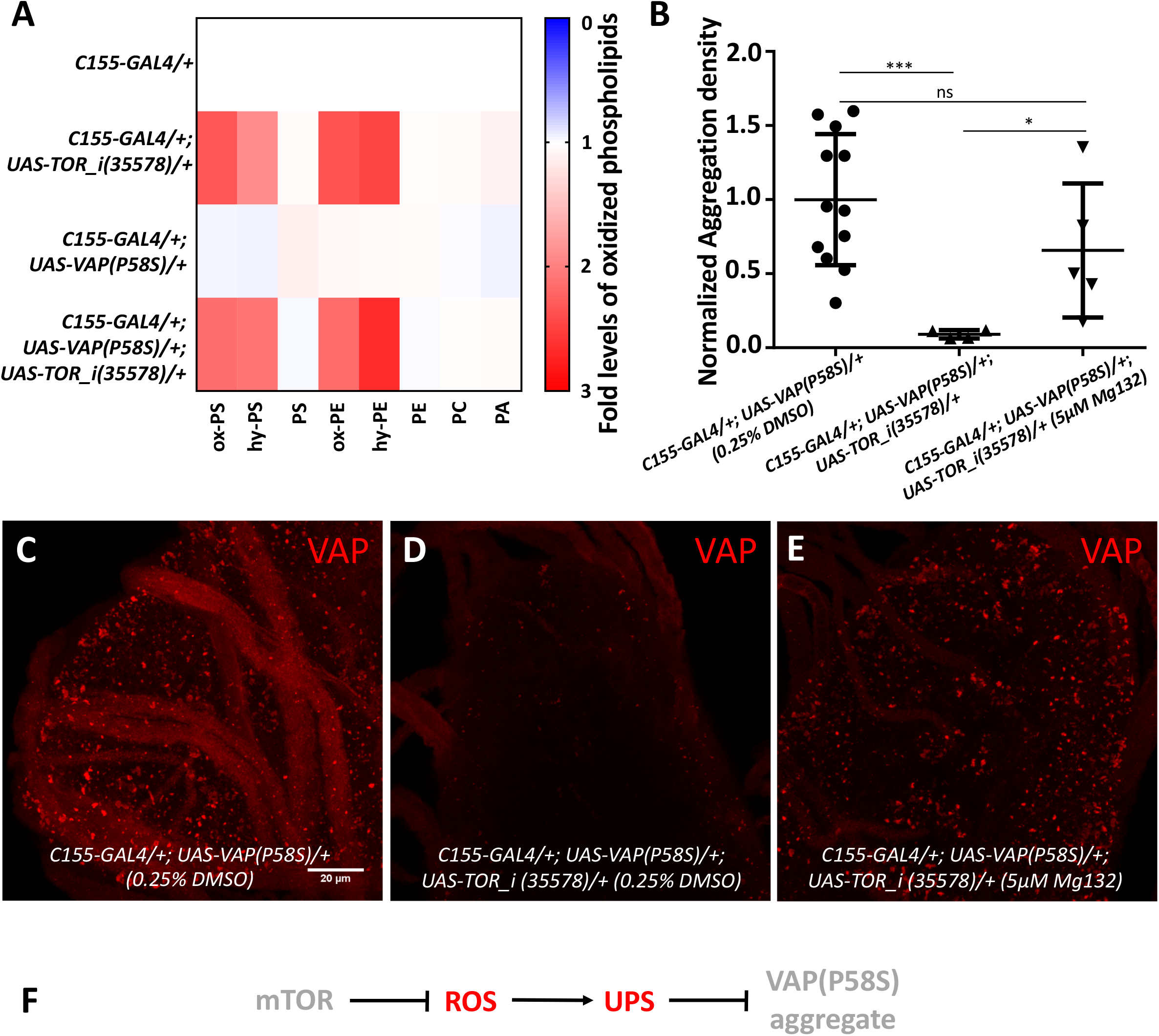
mTOR inhibition increases ROS leading to proteasomal degradation of VAP(P58S) protein/aggregates: **A:** Heat map depicting change in levels of oxidized phospholipids with *TOR* knockdown normalized to *C155-GAL4/+*, quantified using MS in response to ROS generated in third instar larval brains (N=3/4) for the listed genotypes. Statistical tests are described in supplementary table 2 **B:** Plot showing significant decrease in aggregation density in the ventral nerve cord in *C155-GAL4/+; UAS-VAP(P58S); UAS-TOR_i (35578)/+* as compared to *C155-GAL4/+; UAS-VAP(P58S)/+* control. This decrease is partially rescued by feeding 5µM MG132. ANOVA (P-value: **, 0.0042) Fisher’s LSD multiple comparison test (P-values, *<0.05, ***<0.001) **C,D,E:** Representative images of third instar larval brains showing the partial recovery of aggregates upon 5µM MG132 feeding in *C155-GAL4/+; UAS-VAP(P58S)/+; UAS-TOR_i (35578)/+* larvae. All images were taken at the same magnification. **F:** Model depicting the role of mTOR-regulated ROS in activating proteasomal degradation of VAP(P58S) protein/aggregates.

We also explored the possible relationship between *VAP* and ROS at a transcriptional level. Larvae of the control, *CI55-GAL4/+* genotype were hatched and fed on 5mM paraquat, and the brains were dissected at the wandering third instar larval stage. The levels of endogenous *VAP* and *SOD1* mRNA, in response to ROS, were measured using qPCR in control larval brains. We found that endogenous *VAP* mRNA levels were lower in the presence of high levels of ROS (Suppl. Fig. 4E), while *SOD1* mRNA levels remained unchanged (Suppl. Fig. 4F). This result may indicate the presence of a negative feedback loop wherein VAP overexpression leads to accumulation of ROS (Fig. 4C-D), which in turn downregulates endogenous *VAP* transcription.

## Discussion

### A targeted RNAi screen uncovers SOD1, TDP43 and TOR signalling elements as targets to understand dynamics of VAP(P58S) aggregation

*Drosophila* S2R+ cell based whole genome RNAi screens serve as powerful tools due of the relative ease with which transcript knockdown can be achieved (Echeverri and Perrimon 2006). Similar systems have been used for identifying modifiers of aggregation of Huntingtin protein (Zhang *et al*. 2010). Our screen was aimed at enriching genes that are known players in ALS, VAP interactors and proteostasis. First and foremost, we found ALS loci, *SOD1* and *TDP-43* as modifiers of VAP(P58S) aggregation, which we had previously identified as VAP genetic interactors (Deivasigamani *et al*. 2014). In this study, we have explored the interaction between *SOD1* and *VAP*, while *TDP-43* also serves as an exciting candidate for further investigation. TDP-43 has been shown to perturb membrane-associated mitochondrial (Turner *et al*. 2008) sites that are maintained by VAPB-PTPIP51 interactions in mammalian cell culture (Stoica *et al*. 2014). Additionally, TDP-43 proteinopathy has been identified in motor neurons of mice models of VAP(P58S) aggregation (Tudor *et al*. 2010). TDP-43 driven neurodegeneration has also been shown to be modulated by oxidative stress related MAP kinase pathways in a *Drosophila* screen (Zhan *et al*. 2015) and associated with Nrf2 dependent antioxidant pathway (Moujalled *et al*. 2017). In addition to SOD1, we have also identified other ROS related genes such as peroxiredoxin V, NADH dehydrogenase, cytochrome c oxidase, that localise to the mitochondria, perturbation of which will lead to oxidative stress, potentially affecting aggregation kinetics of VAP(P58S).

Secondly, we enriched a subset of targets involved in protein degradation, UPS and autophagy, an *in vivo* validation of which would shed light on the how these aggregates are compartmentalized and managed in the neurons. Thirdly, this screen highlighted specific chaperones that could be involved in the misfolding and formation of VAP(P58S) aggregates providing insight into the initiation of the disease condition. Most importantly, through our previous study (Deivasigamani *et al*. 2014), and our cell-based screen followed by subsequent experimentation, we have established mTOR signalling as a strong modulator of VAP(P58S) aggregation. mTOR signalling responds and integrates signals from nutrients, growth factors, energy, and stress, regulates cellular proteostasis, thus contributing to age-related neurodegenerative diseases (Perluigi *et al*. 2015), making it an attractive target for further investigation in ALS pathogenesis. Indeed, rapamycin, a TORC1 inhibitor, is now being used for phase-II clinical trials for ALS (Mandrioli *et al*. 2018). Lastly, through our screen, targeting processes involved in neurodegeneration, we have identified interactions that point towards a role for VAP as a contributor to a common gene regulatory network (GRN), in agreement with several examples in literature (Tudor *et al*. 2010; van Blitterswijk *et al*. 2012; Prause *et al*. 2013; Deivasigamani *et al*. 2014; Stoica *et al*. 2014; Stoica *et al*. 2016; Paillusson *et al*. 2017). When we compared our list of targets with the results from another fly-based screen for VAP(P58S)-induced eye degeneration (Sanhueza *et al*. 2015), we found no overlap, possibly because of differences in sets of genes screened, cell types, and phenotypes visualized.

### An ROS dependant physiological mechanism that triggers proteasomal clearance of VAP(P58S) aggregation

In our study, we have used a dosage-dependent pan-neuronal GAL4 expression of VAP(P58S) in order to study changes in aggregation in the third instar larval brain. We found two targets, SOD1 and mTOR (Deivasigamani *et al*. 2014), downregulation of which, led to a decrease in VAP(P58S) aggregation accompanied by oxidative stress. We identified a role of ROS in upregulating the proteasomal machinery and, thereby facilitating the degradation of misfolded VAP(P58S) protein/aggregates (Integrated Model; Fig. 8A). However, in absence of ROS, we did not find any change in aggregation density upon pharmacological proteasomal inhibition. This is consistent with the cell culture studies that point towards the downregulation of Ubiquitin-proteasome system (UPS) with VAP(P58S) aggregation as a dominant negative effect on wild type VAP function (Kanekura *et al*. 2006; Gkogkas *et al*. 2008; Papiani *et al*. 2012; Genevini *et al*. 2014). Overexpression of VAP(P58S) or loss of VAP in *Drosophila* has been shown to enhance ER stress in the adult brains and may be a result of suspended proteasomal degradation (Tsuda *et al*. 2008; Moustaqim-Barrette *et al*. 2014). In mice, VAP(P56S) aggregates have been shown to represent an ER-Quality Control (ERQC) compartment that develops as a result of a debilitated ER-Associated Degradative (ERAD) pathway (Kuijpers *et al*. 2013). Indeed, VAP has been shown to interact with UPR sensor AFT6 in mice and the ERAD complex thereby regulating proteostasis and lipid homeostasis in HeLa cell lines (Gkogkas et al, 2008; Ernst et al, 2016). Studies in mammalian cell lines suggest that VAP(P56S) is ubiquitinated, aggregates on the ER membrane and is cleared by the AAA+ valsolin containing protein (VCP)/p97, which interacts with Fas associated factor 1(FAF1) and may use the FFAT motif in FAF1 as an adapter to interact with VAP (Papiani *et al*. 2012; Baron *et al*. 2014). In *Drosophila*, VAP has been shown to be essential for ER homeostasis by maintaining lipid transport, whereas the mutant VAP flies show accumulation of ubiquitinated and membrane proteins in neuronal cells (Moustaqim-Barrette *et al*. 2014). Hence, although ER stress is build up with VAP(P58S) aggregation, it does not lead to subsequent oxidative stress, as shown in our results. This suggests that ROS enhances the proteasomal degradation of VAP(P58S) through an ER stress-independent mechanism. Although neuronal VAP(P58S) aggregates appeared to be non-toxic to flies *per se*, our study highlights the effects of ROS on the dynamics of VAP(P58S) from misfolded protein to aggregate formation and subsequent clearance.

**Figure 8:**
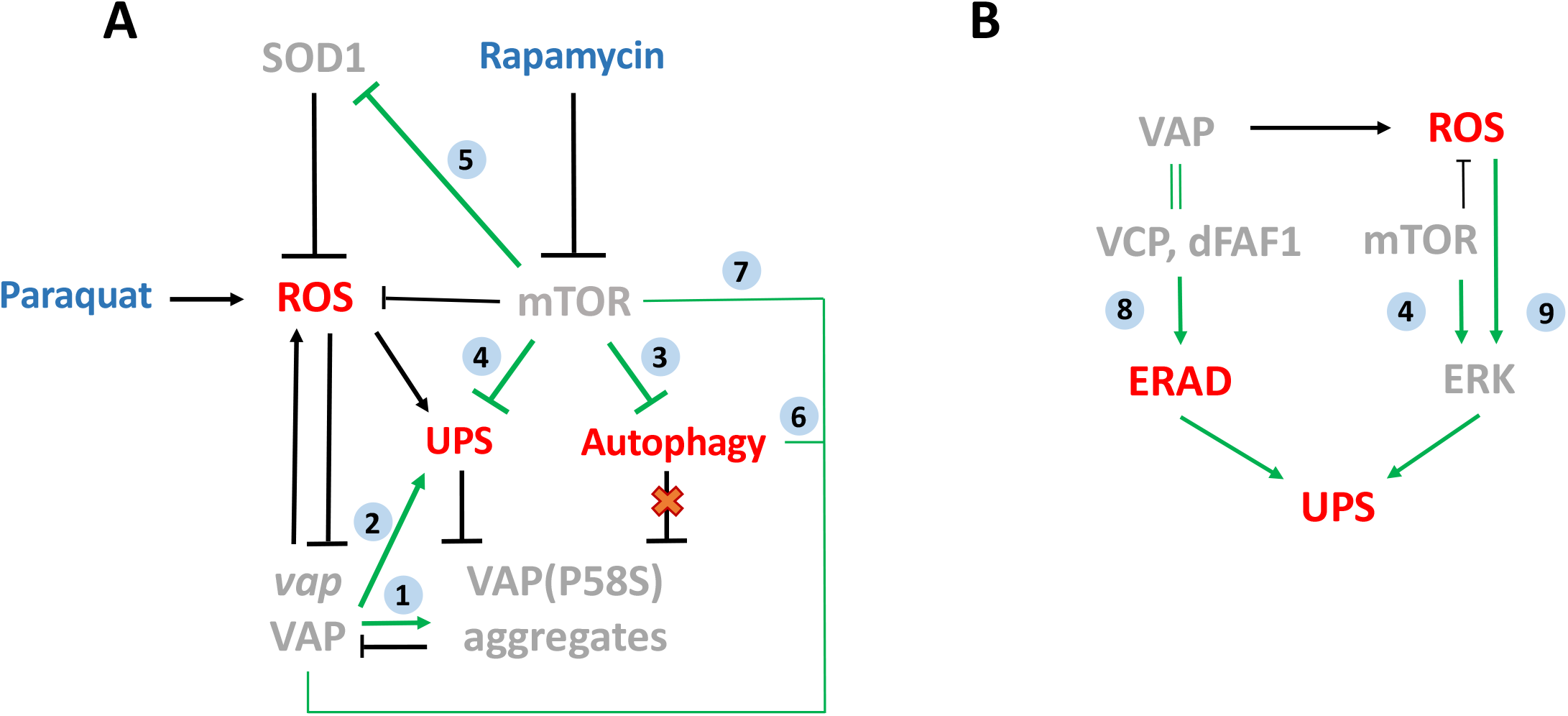
An integrated model for ROS mediated clearance of VAP(P58S) aggregates via UPS. **A:** Model depicting novel relationships of SOD1(ALS1) and mTOR-induced ROS with *vap* and VAP(P58S) aggregates. Clearance of VAP(P58S) protein/aggregates appears to be primarily via the Ubiquitin-Proteosomal system (UPS), triggered by cellular pathways such as mTOR pathway, SOD1 and VAP activity, which in turn regulate ROS. Autophagy does not appear to be a major contributor for aggregate clearance, under the conditions of our experiment. **B:** A hypothetical model proposing the possible link between VAP, ROS and UPS. VAP could regulate the UPS via the ERAD pathway due to its interaction with VCP via dFAF1/Caspar. ROS could be the connecting link between mTOR pathway and ERK pathway that together regulate the components of the proteasomal machinery. The link between VAP and ROS that we have demonstrated could modulate proteasomal activity in the cell. Gray text indicates Genes (*italics*) and proteins (Capitals); Red text indicates cellular mechanisms; Blue text indicates drugs; Arrows: Black: Experimental evidence, this study; Green: Relationship described in literature; Numbers inside blue circles indicate research papers: **1**. Ratnaparkhi *et al*.,2008; **2**. Kanekura *et al*.,2005; Kuijpers *et al*.,2013; **3**. Noda and Ohsumi, 1998; Perluigi *et al*.,2015; **4**. Zhao *et al*.,2015; Rousseau *et al*.,2016; **5**. Sun *et al*., 2012; Tsang *et al*., 2018; **6**. Gomez-Suaga *et al*., 2017; Zhao *et al*., 2018; **7**. Deivaisigamini *et al*., 2014; **8**. Baron *et al*.,2014; Papiani *et al*.,2012; **9**. Cavanaugh *et al*.,2006; Su *et al*.,2014.

### TOR signaling regulates VAP(P58S) dynamics by a UPS dependent and Atg1 independent mechanisms

We previously identified mTOR pathway as a strong regulator of both VAP and VAP(P58S) phenotypes at the neuromuscular junction (Deivasigamani *et al*. 2014). Here, we have shown that inhibition of mTOR pathway also reduces VAP(P58S) aggregation levels in third instar larval brains in presence of ROS. mTOR pathway downregulation is known to activate autophagy (Noda and Ohsumi 1998), a process that has been shown to reduce mutant huntingtin fragments (Ravikumar *et al*. 2004) and amyloid-β levels (Spilman *et al*. 2010) in mice models. Autophagy has been suggested to be upregulated in presence of VAP(P56S) aggregates that also colocalize with the autophagic marker, p62, in mice (Larroquette *et al*. 2015). With *VAP* knockdown in cell culture, autophagy is upregulated due to the loss of calcium homeostasis that arises with the disruption of ER-mitochondrial contact sites (Gomez-Suaga *et al*. 2017a; Gomez-Suaga *et al*. 2017b). However, VAP is also suggested to have a role in autophagosomal biogenesis through direct interaction with autophagy proteins (Zhao *et al*. 2018). In our study, we do not observe any clearance of VAP(P58S) aggregation with activation of Atg1, indicating that clearance observed with mTOR inhibition may be an effect of one or more of its downstream processes (Fig. 8A).

mTOR and SOD1 have been shown to be genetic interactors in *Drosophila* with mTOR inhibition enhancing the lifespan defect incurred with SOD1 knockdown (SUN *et al*. 2012). Recently, mTOR has been directly shown to regulate SOD1 activity by its phosphorylation based on nutrient availability in yeast and mammalian cells (Tsang *et al*. 2018). Although this phosphorylation site does not appear to be conserved in *Drosophila*, this study demonstrates the role of mTOR pathway in regulating ROS via SOD1. mTOR inhibition, specifically, mTORC1 has also been shown to activate proteasomal degradation independent of its other targets, such as, 4EBP, S6K and Ulk (Cavanaugh *et al*. 2006; Zhao *et al*. 2015). An evolutionarily conserved regulation of components of proteasomal assembly by mTORC1 via Mpk1/ERK5 has been reported in yeast as well as mammalian cell culture (Rousseau and Bertolotti 2016). ERK5 signalling has been implicated in neuroprotective roles in response to mild levels of oxidative stress (Cavanaugh *et al*. 2006; Su *et al*. 2014). These studies suggest that ROS regulation by mTOR inhibition via SOD1 and ERK5, serves as a plausible mechanism for the proteasomal degradation of VAP(P58S) protein/aggregation, and by extension, the rescue of VAP(P58S) NMJ phenotype (Deivasigamani *et al*. 2014) (Fig. 8B).

### Increase in ROS by VAP, but not VAP(P58S) expression

SOD1-associated elevation in ROS levels and oxidative stress is suggested as a plausible factor of motor neuron death in ALS (Barber *et al*. 2006; Saccon *et al*. 2013). Teuling *et al*., 2007 (Teuling *et al*. 2007) have shown that VAPB protein levels decrease in an age-dependent manner in a mouse model of SOD1-G93A, providing the first evidence of a link between *ALS1* and *VAP/ALS8*. We now find that overexpressed VAP, unlike VAP(P58S), promotes the accumulation of ROS in the system. This is consistent with a study that shows lowered ROS in a *vpr* (VAP ortholog) mutant of *C. elegans* in response to increased mitochondrial connectivity and altered function (Han *et al*. 2012). VAP neuronal overexpression in *Drosophila* has also been shown to increase bouton number (Pennetta *et al*. 2002) similar to SOD1 mutant phenotype at the NMJ (Milton *et al*. 2011), and is correlated with increased ROS in both scenarios. VAP may be important in regulating pathways that respond to changes in ROS levels, such as mTOR and ERK pathways that can regulate UPS (Rousseau and Bertolotti 2016). VAP also modulates ERAD (and UPS), via its interaction with VCP and FAF1 (Papiani *et al*. 2012; BARON *et al*. 2014). We hypothesize that the interaction between VAP and ROS could lead to crosstalk between these pathways regulating global proteostasis (Fig. 8B).

### ROS may regulate VAP levels by regulating VAP transcription

In our study, we have found that in presence of ROS, *VAP* transcription is downregulated in wild type flies. We had previously shown that *SOD1* knockdown rescues VAP macrochaetae phenotype (Deivasigamani *et al*. 2014), which may be a consequence of excessive ROS accumulation, and subsequent downregulation of VAP levels and function. Two independent studies (Qiu *et al*. 2013; Kim *et al*. 2016), that overexpressed VAPB in *ALS1* (SOD1-G93A) mice as an attempt at rescuing ALS defects, found contradictory observations, owing mainly to differences in expression levels of the protein. *VAPB* mRNA levels are known to be lowered in spinal cords of patients with sporadic ALS (Anagnostou *et al*. 2010), as well as in IPSC-derived motor neurons from ALS8 patients (Mitne-Neto *et al*. 2007). It has also been reported that VAPB staining in motor neurons of sporadic patients is increased showing “punctate accumulation” that colocalize with early endosomal marker, Rab5 (Sanhueza *et al*. 2015). Based on our results and taking into consideration earlier observations (Teuling *et al*. 2007; Anagnostou *et al*. 2010; Deivasigamani *et al*. 2014), we submit that in the ALS disease scenario, increased VAP accumulates ROS that initiates a negative feedback loop resulting in downregulation of VAP, at the transcript level (Fig. 8A). It remains to be tested whether ROS-activated pathways such as MAP kinase pathways or mTOR pathway, could directly control VAP expression. This VAP/ROS regulation that we have uncovered may have significant implications in ALS pathogenesis for both sporadic and familial ALS.

In Summary, we find that the dynamics of VAP(P58S) neural aggregates, a species intimately linked to disease in the human context, is sensitive to levels of ROS. Change in physiological levels of ROS appear to dictate the equilibrium between the aggregated and non-aggregated forms. The cellular levels of ROS are themselves dictated by well characterized regulatory mechanisms that include ROS generators and scavengers. As shown in this study, TOR signalling and VAP/VAP(P58S) expression levels would contribute to the extent of aggregation, and may act as regulatory feedback loops to regulate physiological ROS levels. SOD1, VAP/ALS8, TOR and ROS are part of physiological regulatory circuit that maintains levels of VAP(P58S) aggregates.

## Materials & Methods

### Generation of constructs and dsRNA

The cDNA sequence of VAP and VAP(P58S) mutant were cloned into *pRM-GFP* plasmid (Bhaskar *et al*. 2000) to generate both N and C-terminal GFP fusions, using the *EcoR1* restriction site. The pRM-GFP vector has GFP cloned into pRM-HA3 vector at the *BamHI* site. 500 uM CuSO4 was used to drive expression is S2R+ cells after transient transfections. dsRNA for the secondary screen was generated using MEGAscript® T7 Kit (AM1333) by ThermoFisher Scientific. Template for dsRNA was generated by using cDNA as template, prepared from flies. Primers for the same were ordered from Sigma.

### Handling of Schneider cells

*Drosophila* S2R+ cells were maintained in Schneider cell Media (#21720–024; GIBCO) with 10% Heat inactivated Fetal Bovine Serum (FBS, #10270; GIBCO). Batches of cells were frozen in 10% DMSO (D2650; Sigma) and stored in liquid nitrogen following DRSC protocol (http://www.flyrnai.org/DRSC-PRC.html). In general, after reviving, cells were discarded after 25–30 passages. Cells were maintained at 23° C, and split every 4 days at a ratio of 1:5.

### Cell culture and generation of S2R+ stable lines

Stable S2R+ cell lines were generated by co-transfecting with pRM-HA3 constructs of VAP:GFP, VAP(P58S):GFP or GFP along with pCo-Hygro in 20:1 ratio, using Effectene (QIAGEN) and/or Mirus TransIT 2020 (MIR 5400), and selected under 250 µg/ml of hygromycin (Sigma) for 10-15 passages. Stable as well as transiently transfected cell lines were induced to express the gene of interest under a metallothionein promoter using increasing concentrations 250μM, 500μM, 750μM and 1000μM of copper sulphate and analysed at 12, 24, 36 and 48 hours post induction. Transient transfections assays were performed using Mirus TransIT-2020 (MIR 5400) transfection reagent. Protocol for dsRNA knockdown assay was modified from (Rogers and Rogers 2008). Fixation, DAPI staining and imaging was done using EVOS FL Auto Cell Imaging system. Super-resolution images of fixed VAP:GFP and VAP(P58S):GFP cells were acquired using Leica SR GSD 3D system.

### Western blotting

Cells were centrifuged at 3000 rpm for 5 minutes in Eppendorf 5414R centrifuge. The pellet was resuspended in 20 μl of supernatant and boiled with 1X SDS Dye at 95°C. Samples were centrifuged again at 10000 rcf for 10 minutes. Cell extracts were separated by 12% SDS-PAGE and transferred onto 0.45 μm PVDF membrane (Millipore). Membranes were blocked for 1 hour in 5% skimmed milk in 1X TBS containing 0.1% Tween-20 at room temperature and probed with 1:10,000 diluted mouse anti-Tubulin (T6074; Sigma-Aldrich) and 1:5,000 diluted mouse anti-GFP (Roche life science), overnight at 4°c (12 hours). Anti-rabbit and anti-mouse secondary antibodies conjugated to horseradish peroxide (Pierce) were used at a dilution of 1:10,000 for 1 hour at room temperature. Blots were developed with Immobilon Chemiluminescent Substrate (LuminataClassico Western HRP substrate from Millipore) using a LAS4000 Fuji imaging System.

### GO analysis

The list of genes and Gene Ontology (GO) information was obtained based on Flybase (http://flybase.org) (Marygold *et al*. 2013) entries. Genes were categorized manually in the broad categories of ALS genes, VAP interactome (Deivasigamani *et al*. 2014) and proteostasis. List of ALS loci and ALS related genes were obtained from http://alsod.iop.kcl.ac.uk/ (Wroe *et al*. 2008). The *Drosophila melanogaster* homologs of these ALS genes were identified using Ensembl biomart tool (http://asia.ensembl.org/biomart/martview) and Flybase batch download tool. Human orthologs of the target genes listed in Suppl. Table 1C and 1D were identified using DRSC Integrative Ortholog Prediction Tool (DIOPT) (http://www.flyrnai.org/cgibin/DRSC_orthologs.pl).

### High through-put screen, and image acquisition

The screen was performed at the screening facility at CCAMP-NCBS, Bangalore (http://ccamp.res.in/HTS-HCI). dsRNA for the high throughput screen was generated and plated into sixteen 384 well plates by Chromous Biotech, Bangalore in preparation for the experiment. The library used as a template for generating dsRNAs was procured from Open Biosystems (RDM1189 and RDM4220). 50 μl of cells (3 × 10^6 / ml) were plated in each well for the 384 well flat bottom plates obtained from Corning. Each target dsRNA knockdown experiment was done in triplicate, randomly arranged in the 384 well plate. The cells were treated with 10 μg/ml of dsRNA for 48 hours, followed by induction with 500 μM CuSO_4_. The cells were fixed and imaged at 24 and 36 hours post CuSO_4_ induction. Fixation was done with 4% PFA in 1X PBS, washed twice with 1X PBS, treated with 0.05µg/ml DAPI and followed with two washes with 1X PBS. Each plate contained 7 negative controls occupying 42 wells. 114 unique genes were screened in each plate. Few genes were kept as overlap between multiple plates to check for their consistency and reproducibility. Imaging for the high throughput screen was performed by THERMO Array Scan VTI HCS system. Dual-channel images from ten fields in each well were captured using a 20X air objective and an EMCCD camera. The FITC (488nm) channel was used for imaging VAP(P58S):GFP aggregates and the DAPI (405nm) channel for imaging cell nuclei. 10 fields were imaged in each well and around 400 cells were imaged per field. In well triplicates, around 12,000 cells were imaged for each dsRNA knockdown.

### High throughput data analysis

Images from the FITC and DAPI channels in each site were read using the Bio-Formats MATLAB toolbox (Linkert *et al*. 2010) and were processed using custom MATLAB scripts. The segmentation was done using the DAPI images and the extraction of pixel intensities was done on the FITC channel. Illumination correction was performed as a pre-processing step on the DAPI Images and individual nuclei were segmented after a contrast stretching routine was applied. The identified objects were further filtered for outliers, based on a size-based cutoffs and the individual 8-connected components were labelled as separate nuclei. Under 20x magnification we estimated the cellular radius to be around 10 pixels corresponding to 5 μm. Thus, labelled cellular objects (ROIs), were obtained by dilating the centroids of each nuclei by 10 pixels. Around 400 ROIs were obtained from each field consistent with manually counted cells in these images. The resultant ROI’s were further filtered for clumps and out of focus objects. The GFP intensities were obtained for these ROI’s post a local background correction of the FITC images (with a disk size of 3 pixels). Average and total intensities were calculated from the pixel data obtained from every cell/ROI from these FITC images. A Kolmogorov-Smirnov-like (KS) statistic was used to assign Z-scores to each gene on plate as reported by (Dey *et al*. 2014). A statistically significant threshold was obtained for the triplicate data using monte-carlo simulations. Genes were classified as hits, if it occurred two or more times above a given Z-score threshold. The false positive rates for both parameters at both time points was zero. The false negative rates for average intensity for 24 hours-time point was 0.2523 and for 36 hours-time point was 0.361. The false negative rates for total intensity for 24 hours-time point was 0.3838 and for 36 hours-time point was 0.3164.

### Fly husbandry and brain aggregation assay

Fly lines were maintained on standard corn meal agar medium. *UAS-GAL4* system was used for overexpression of transgenes. *UAS-VAP* wildtype, *UAS-VAP(P58S)* and *C155-GAL4* lines used for fly experiments have been described earlier (Ratnaparkhi *et al*. 2008; Deivasigamani *et al*. 2014). Canton S flies were used as wildtype control. *UAS-VAP_i* (27312), *UAS-SOD1_i*(34616, 29389, 36804) and *UAS-TOR_i* (35578) where the suffix ‘I’ indicates an RNAi line, and *UAS-SOD1* (24750, 33605) were obtained from BDSC. Clone for UAS-FLAG-HA tagged SOD1 in pUASt vector was obtained for expression in *Drosophila* from DGRC and injected in the NCBS-CCAMP transgenic facility. *UAS-Atg1* line was kindly provided by Dr. Chen, Academia Sinica; the line was validated in the wing using *ptc-GAL4* as described (Chen *et al*. 2008). Experimental Crosses were set at 18°C, 25°C or 28°C, as indicated. Brains were dissected from third instar larvae and processed for immunostaining assay. 4% paraformaldehyde containing 0.1% Triton-X was used for fixation followed by washes with 1X PBS. Blocking treatment and washes were performed with 0.3% Triton-X with 2% BSA. Brains were stained with 1:500 diluted anti-VAP antibody and 1:1000 anti-rabbit secondary (Invitrogen) was used. Z-stacks of five-ten brains for each sample were imaged under 63X oil objective of Ziess LSM 710 Confocal Microscope. The number of aggregates were quantified per cubic micron of the ventral nerve cord, defined as “aggregation density” using the Huygen professional software. The high intensity puncta were considered as aggregates. An arbitrary threshold was set for controls as well as for test samples that achieved removing low intensity background signal emitted by the tissue, along with separation of high intensity puncta that were adjacent to one another. An object filter was used to remove objects of size greater than 1000 pixels and garbage size smaller than 10 pixels was excluded. Three 3D region of interests of fixed size were drawn along the tip of the ventral nerve cord and the number of aggregates were counted from each of these ROIs and averaged for each animal. The volume (in cubic micron) of ROI depicting the thickness of the brain tissue was measured as the range of the z-stack of the image. The aggregation density obtained for each brain has been normalised to the mean of the control group, *C155-GAL4; UAS-VAP(P58S)* (+ 0.25% DMSO, in case of DMSO-soluble drug experiments) and plotted as “normalized aggregation density” in each graph. Student t-test and one-way ANOVA were used to measure statistical significance.

### Drug treatment

Cells were exposed to 10mM and 20mM Paraquat dichloride hydrate (500mM, 36541-Sigma-aldrich) for 24 hours prior to protein induction with 500µM copper sulphate. Fixation, DAPI staining and imaging was done using EVOS FL Auto Cell Imaging system. For flies, 10-12 virgins were placed with CS males, for each genotype and animals were allowed to mate for 24 hours and transferred to standard cornmeal fly media containing paraquat (0.05mM, 0.5mM, and 5mM), MG132 (5µM), rapamycin (200nM) or DMSO (0.25%).

### Oxyblot assay

Third instar larval brains were lysed in RIPA containing 50 mM DTT and centrifuged at 10000 rcf. The lysate containing 10µg of protein was incubated with 2,4-dinitrophenylhydrazine (DNPH) to derivatize the carbonyl groups of oxidized proteins with 2,4-dinitrophenylhydrazone (DNP-hydrazone) as described by the Oxyblot Protein Oxidation Detection Kit (S7150) from EMD Milipore. The derivatized protein lysate was separated on a 12% SDS-PAGE and transferred onto 0.45 μm PVDF membrane (Millipore). Oxidized protein levels in the lysate were detected by probing with anti-DNP antibody on western blot as per the Oxyblot Protein Oxidation Detection Kit manual.

### Lipid extraction and targeted LC-MS lipidomics

All MS quantitation phospholipid standards were purchased from Avanti Polar Lipids Inc., USA. The brain samples were washed with PBS (x 3 times), and transferred into a glass vial using 1 mL PBS. 3 mL of 2:1 (vol/vol) CHCl_3_: MeOH with the internal standard mix (1 nmol 17:1 FFA, 100 pmol each of 17:0-20:4 PS, 17:0-20:4 PC, 17:0-20:4 PE, and 17:0-20:4 PA) was added, and the mixture was vigorously vortexed. The two phases were separated by centrifugation at 2800 × g for 5 minutes. The organic phase (bottom) was removed, 50 μL of formic acid was added to acidify the aqueous homogenate (to enhance extraction of phospholipids), and CHCl_3_ was added to make up 4 mL volume. The mixture was vortexed, and separated using centrifugation described above. Both the organic extracts were pooled, and dried under a stream of N_2_. The lipidome was re-solubilized in 200 μL of 2:1 (vol/vol) CHCl_3_: MeOH, and 20 μL was used for the targeted LC-MS analysis (Kamat *et al*. 2015). All the phospholipid species analyzed in this study were quantified using the multiple reaction monitoring (MRM) method on an AbSciex QTrap 4500 LC-MS with a Shimadzu Exion-LC series quaternary pump. All data was collected using the Acquisition mode of the Analyst software, and analyzed using the Quantitate mode of the same software. The LC separation was achieved using a Gemini 5U C-18 column (Phenomenex, 5 μm, 50 × 4.6 mm) coupled to a Gemini guard column (Phenomenex, 4 × 3 mm, Phenomenex security cartridge). The LC solvents were: For positive mode: buffer A: 95:5 (vol/vol) H_2_O: MeOH + 0.1% formic acid + 10 mM ammonium formate; and buffer B: 60:35:5 (vol/vol) iPrOH: MeOH: H_2_O + 0.1% formic acid + 10 mM ammonium formate, For Negative mode: buffer A: 95:5 (vol/vol) H_2_O: MeOH + 0.1% ammonium hydroxide; and buffer B: 60:35:5 (vol/vol) iPrOH: MeOH: H_2_O + 0.1% ammonium hydroxide. All the MS based lipid estimations was performed using an electrospray ion source, using the following MS parameters: ion source = turbo spray, collision gas = medium, curtain gas = 20 L/min, ion spray voltage = 4500 V, temperature = 400°C. A typical LC-run consisted of 55 minutes, with the following solvent run sequence post injection: 0.3 ml/min 0% buffer B for 5 minutes, 0.5 ml/min 0% buffer B for 5 minutes, 0.5 ml/min linear gradient of buffer B from 0-100% over 25 minutes, 0.5 ml/min of 100% buffer B for 10 minutes, and re-equilibration with 0.5 ml/min of 0% buffer B for 10 minutes. A detailed list of all the species targeted in this MRM study, describing the precursor parent ion mass and adduct, the product ion targeted can be found in Suppl. Table 2. All the endogenous lipid species were quantified by measuring the area under the curve in comparison to the respective internal standard, and then normalizing to the number of larval brains. All oxidized phospholipids detected were normalized to the corresponding unoxidized phospholipid internal standard. All the data is represented as mean ± s. e. m. of 4 biological replicates per genotype.

### mRNA isolation, cDNA preparation and qRT PCR

About 1 µg of mRNA was isolated from 12-18 third instar larval brains using Direct-zol™ RNA MicroPrep Kit (R2062) from Zymo Research. The cDNA reaction was carried out using High Capacity cDNA Reverse Transcriptase Kit (4368814) by Applied Biosystems. The qPCR reaction was carried out using KAPA SYBR FAST (KK4602) by Sigma using Replex Mastercycler by Eppendorf. The experiment was carried out in three biological replicates with technical triplicates.

## Acknowledgements

The S2R+ screen was carried out as a paid service at the NCBS:C-CAMP high throughput screening facility. At NCBS, we thank Dr. Satyajit Mayor for his support; MS Shahab Uddin, Lokavya Kurup and Vandana for technical assistance during the execution of the screen; Kausik Chakraborty, IGIB for advice on the analysis of the screen. We thank Bloomington Drosophila Stock Center (BDSC), Indiana, supported by NIH grant P40OD018537, for fly stocks; Drosophila Genome Research Centre (DGRC), Indiana supported by NIH grant 2P40OD010949 for vectors and clones; TRiP collection at Harvard Medical School (NIH/NIGMS R01-GM084947) for providing transgenic RNAi fly stocks. We thank IISER Microscopy/Confocal Facility and Dr. Nagaraj Balasubramaniam for access to the EVOS system. Shubham Singh and Shabnam Patil are thanked for technical assistance. This work is funded by a research grant from the Department of Biotechnology, Govt. of India (BT/PR8636/AGR/36/786/2013) and Department of Science and Technology, Science and Engineering Research Board (DST-SERB), Govt. of India (EMR/2014/000367) to GR, a DST-SERB Early Career Research Award in Life Sciences (ECR/2016/001261) to SSK, and a DST-FIST infrastructure development grant to the IISER Pune Biology Department. LP was a UG student at IISER and carried out the S2R+ screen at NCBS. KC and SD are/were graduate students supported by research fellowships from CSIR, Govt. of India. KC is an awardee of the DMM conference travel grant. We thank; Anuradha Ratnaparkhi for discussions and comments on the Manuscript, Richa Rikhy for helpful discussions.

## Author Contributions

GR conceived the project and designed the experiments, with input from KC, LP, SSK and SD. KC and LP performed all the experiments. BR wrote the MATLAB code to analyse the screen. SSK contributed by designing and overseeing experiments related to oxidation of proteins and lipids. GR, KC, LP, BR, SD and SSK analysed the data and wrote the Manuscript. The authors declare no conflict of interest.

## Supplementary Material

**Suppl. Figures along with their legends are part of the Main Manuscript. Suppl. Tables (described below) have been uploaded formally as ‘Suppl. Files’**.

**Table 1 (Suppl. Table1.xls).** **A**. List of 900 genes utilized for the screen. List is sorted alphabetically based on gene symbol. **B**. 900 genes, utilized for the screen, classified and listed into 10 categories associated with ALS or VAP or proteostasis. **C**. List of 150 modifiers of VAP(P58S) aggregation, based on average cell intensity, along with their human orthologs. **D**. List of 85 modifiers of VAP(P58S) aggregation, based on total cell intensity, along with their human orthologs.

**Table 2 (Suppl. Table2.xls).** **A**. Details of the MRM transitions for the different phospholipids measured **B**. LC-MS quantitation of the different phospholipids for different genotypes and paraquat treatment. **C**. LC-MS quantitation of the different phospholipids for knockdown of *TOR*.

**SUPPLEMENTARY FIGURE 1:**
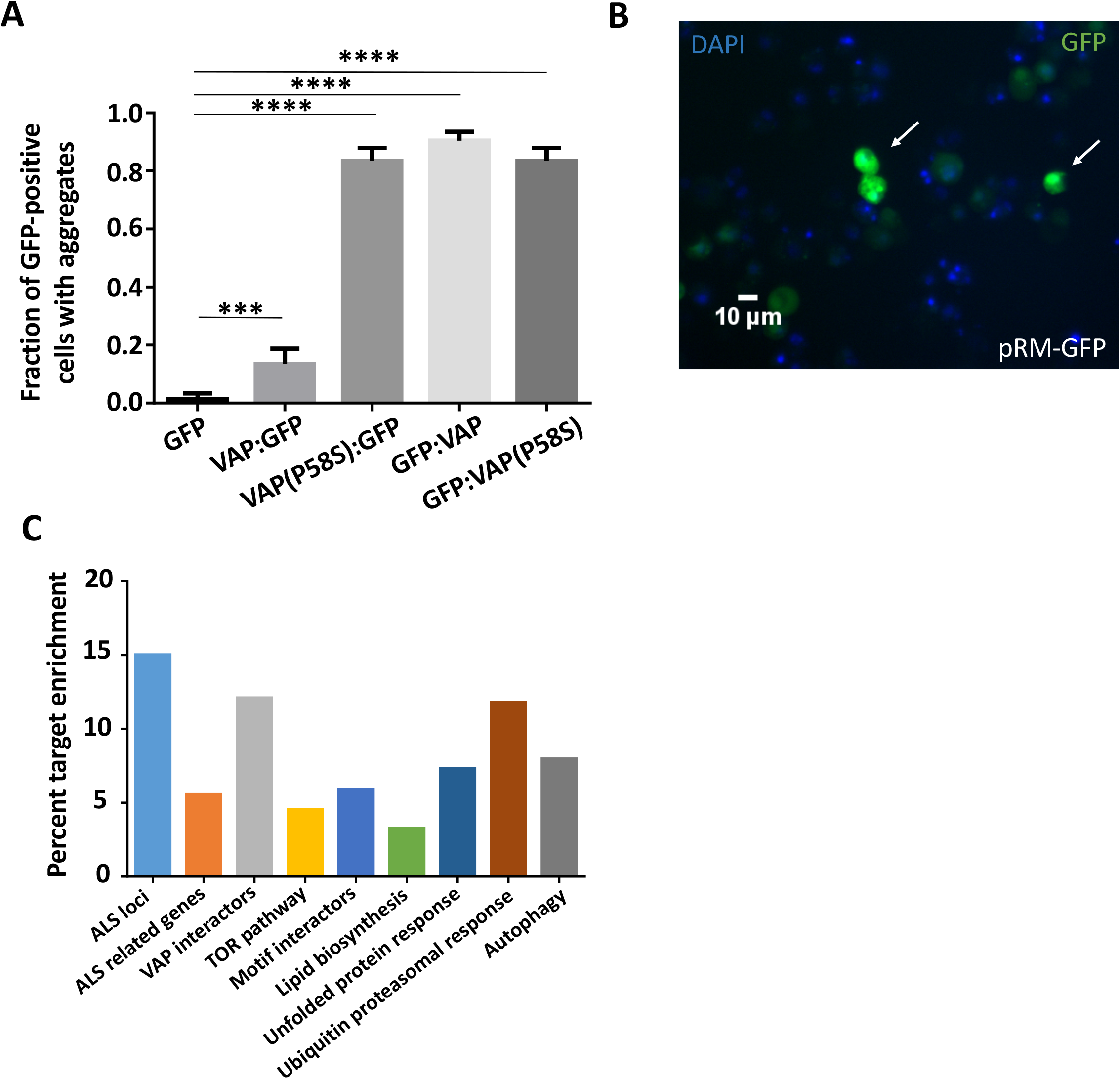
**A:** Fraction of GFP-positive cells showing aggregates plotted for transiently transfected with C-terminally or N-terminally tagged GFP constructs of VAP or VAP(P58S) and only CFP construct at 24 hours post 500µM CuSO_4_ induction. Unlike C-terminally tagged VAP, N-terminally tagged VAP forms aggregates as compared to GFP alone. Both C and N-terminally tagged VAP(P58S) proteins form aggregates. ANOVA (P-value: ****<0.0001) Fisher’s LSD multiple comparison test (P-values, ***<0.001, ****<0.0001). **B:** Homogenous cytoplasmic expression of GFP in S2R+ cells. **C:** The end result of the screen: a list of 85 genes identified based on total cell intensity as a parameter; these genes are predicted to modify aggregation of VAP(P58S):GFP. Graph displays the percent fold enrichment of targets within each gene category. Genes are listed in *Suppl. Table 1D*.

**SUPPLEMENTARY FIGURE 2:**
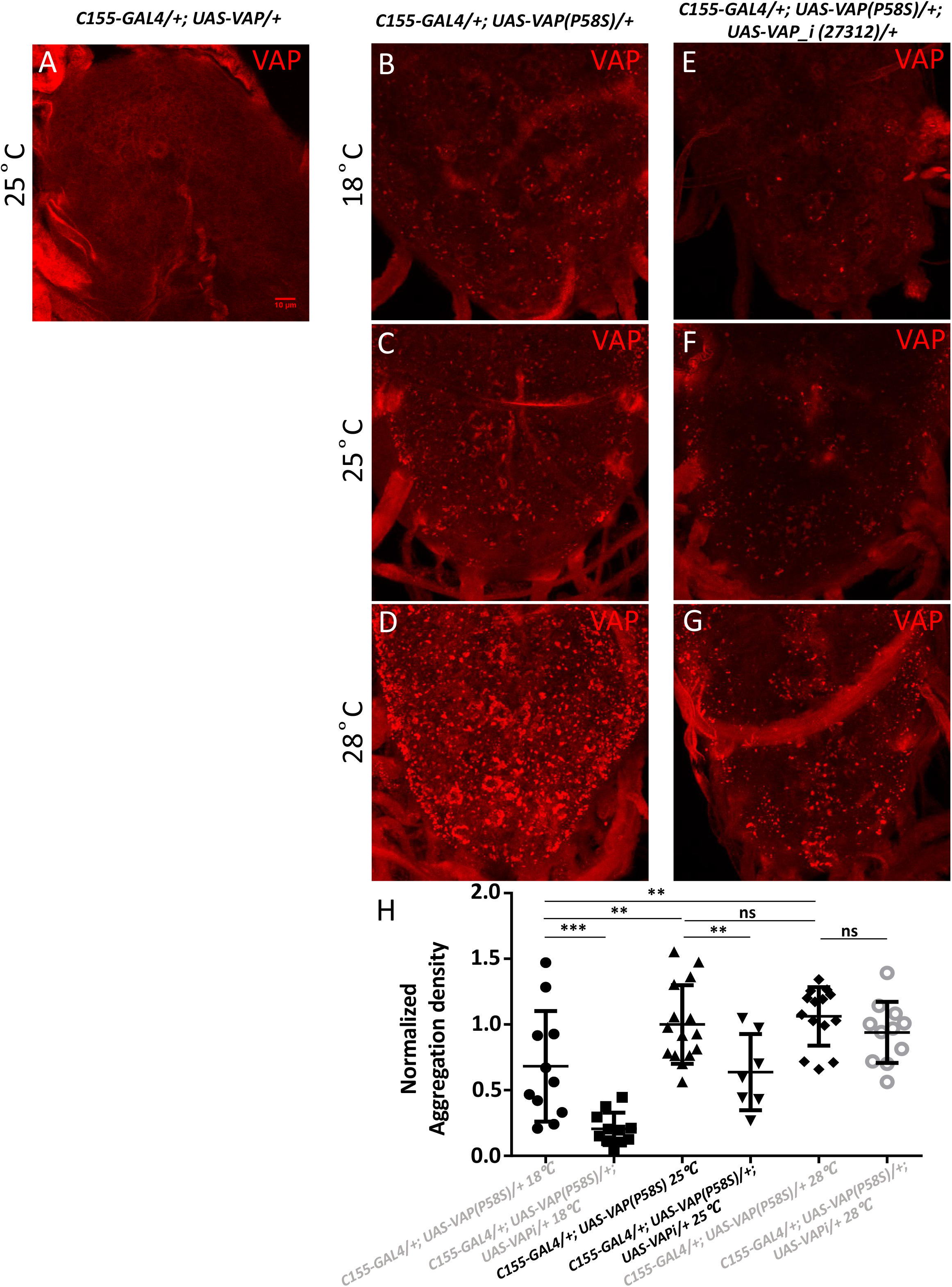
A system for measuring VAP(P58S) aggregation in the larval brain. **A:** Overexpression of VAP in the ventral nerve cord of the third instar larval brain, driven by pan-neuronal *C155-GAL4*, immunostained with rabbit anti-CCD (VAP) antibody, shows membrane localization. **B-D**: Overexpression of VAP(P58S) is visualized as inclusions in the third instar larval brains. Temperature dependent increase in aggregation density is seen in the ventral nerve cord in *C155-GAL4/+; UAS-VAP(P58S)/+* larvae. **E-G**: Knockdown of *VAP* in *C155-GAL4/+; UAS-VAP(P58S)/+* larvae leads to a corresponding decrease in aggregation density at each temperature. **H**: Plot showing significant increase in VAP(P58S) aggregation density with increase in temperature, and a significant decrease in aggregation density in the ventral nerve cord in *C155-GAL4/+; UAS-VAP(P58S); UASVAP_i (27312)/+* as compared to *C155-GAL4/+; UAS-VAP(P58S)/+* control in a temperature dependent manner. All images were taken at the same magnification. ANOVA (P-value: ****<0.0001)Fisher’s LSD multiple comparison test (P values, *<0.05, **<0.01, ***<0.001).

**SUPPLEMENTARY FIGURE 3:**
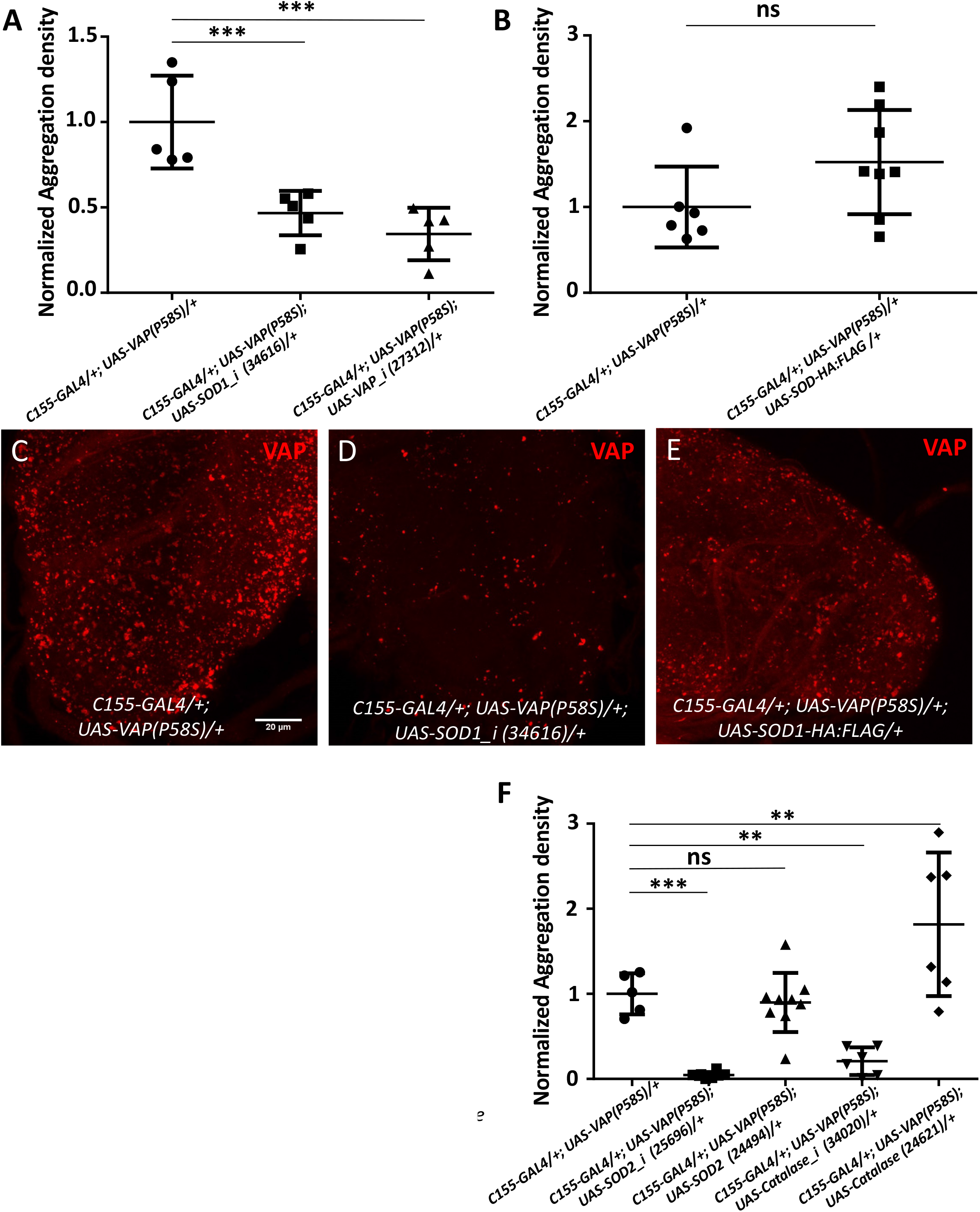
ROS scavenging genes modulate VAP(P58S) aggregation density in the third instar larval brain. **A:** *SOD1* knockdown decreases aggregation density. ANOVA (P-value ***, 0.0004) Fisher’s LSD multiple comparison test (P-value, ***<0.001) **B:** *SOD1:HA:Flag* overexpression does not affect aggregation density. Student’s t test (P-value: 0.1066) **C, D, E:** Representative images of the ventral nerve cord showing aggregation of VAP(P58S) **(C)**, with *SOD1* knockdown **(D)**, and with *SOD1-HA:Flag* overexpression **(E)**. All images were taken at the same magnification. **F:** *SOD2* or *Catalase* knockdown reduces aggregation density. Overexpression of *SOD2* does not change aggregation density, however overexpression of *Catalase* increases aggregation density. The ‘*_i*’ appended to the gene name indicates an RNAi line. ANOVA (P-value: ****<0.0001) Fisher’s LSD multiple comparison test (P-value, **<0.01, ***<0.001).

**SUPPLEMENTARY FIGURE 4:**
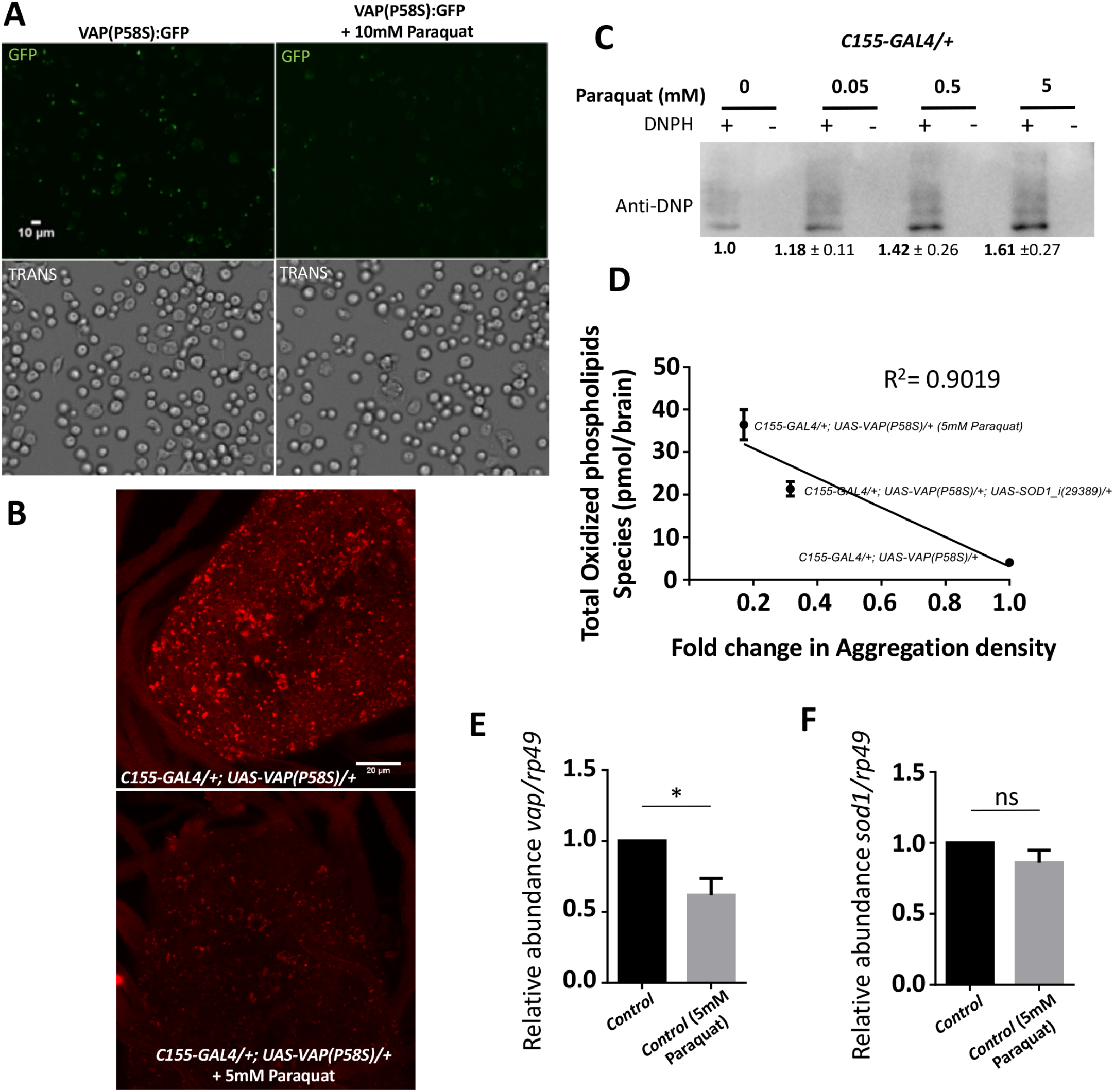
ROS levels are modulated by SOD1 and VAP and vice-versa. **A:** 10 mM Paraquat treatment for 4 hour, prior to inducing VAP(P58S):GFP in stable S2R+ cell line, reduces the fraction of cells showing aggregation observed 24 hours post-induction. **B:** Feeding 5 mM paraquat decreases aggregation density in the ventral nerve cord of third instar larval brains in *C155-GAL4/+; UAS-VAP(P58S)/+* flies. All images are taken at the same magnification. **C:** Higher levels of protein oxidation in larval brains (N=10) is seen using Oxyblot, in response to Paraquat feeding. This experiment serves as a calibration/standard for Fig. 4D. Values below the gel indicate fold intensity of the strongest band, when compared to the control (0 mM papraquat). **D:** Inverse correlation between total oxidized phospholipids and fold change in aggregation density. **E:** Relative mRNA levels *VAP*, in the larval brain are lowered on treatment with 5mM paraquat suggesting that high levels of ROS may negatively regulate *VAP* transcripts. Student's t test was performed. P-value *<0.05 **F:** Relative mRNA levels of *sodl*, in the larval brain, do not change in larvae fed with 5mM paraquat.

